# Spatiotemporal Modeling of Host-Pathogen Interactions using Level Set Method

**DOI:** 10.1101/2024.08.09.607313

**Authors:** Sheila Rae Permanes, Melen Leclerc, Youcef Mammeri

## Abstract

Phenotyping host-pathogen interactions is crucial for understanding infectious diseases in plants. Traditionally, this process has relied on visual assessments or manual measurements, which can be subjective and labor-intensive. Recent advances in image processing and mathematical modeling enable precise and high-throughput phenotyping. In this study, we propose an innovative approach in plant pathology by combining image processing techniques with the level set method. This integrated approach leverages the strengths of both methodologies to provide accurate, robust, and detailed analysis of leaf and lesion evolution. By employing this combination, we achieve precise delineation of lesion boundaries and track their progression over time, offering clear visual feedback. This enhances the ability of the method to monitor plant health status comprehensively. The results, which track the growth of *Peyronellaea pinodes* on the stipules of two pea cultivars and the associated leaf deformation, provide an accurate visual representation of disease progression. This model represents a significant advancement in plant disease phenotyping, offering precise and detailed insights that can enhance our understanding of host-pathogen interactions.

## 1. Introduction

Phenotyping host-pathogen interactions is central for the study of infectious diseases. In plant pathology, it generally involves assessing quantitative traits of aggressiveness (pathogen point of view) or resistance (host point of view). These traits describe the main stages of pathogens life-cycle into host plants and are used to understand the adaptation of pathogens to plant resistance and to identify quantitative trait loci for both host resistance and pathogen aggressiveness (Pariaud et al., 2009; Lannou, 2012). This is particularly important for supporting plant breeding for resistance, as the use of resistant cultivars remains the best alternative to pesticides in agriculture. Practically, pathologists often use plant inoculations in controlled conditions and assess disease development. Historically, the phenotyping of plant-pathogen interactions relied on visual rating or manual measurements. The recent development of imaging techniques and computer vision methods for plant phenotyping now enables automatic, high-precision and high-throughput phenotyping (Bock et al., 2020; Mahlein, 2016).

The use of mathematical epidemiological models is known to offer means for analyzing empirical data, estimating key quantitative traits of the pathogen and better understanding how plant resistance impacts its life-cycle into host tissues. This is especially true for pathogens that induce growing lesions for which the proper estimation of pathogen traits is not necessarily trivial and require a longitudinal monitoring and modelling (Leclerc et al., 2019). The work of Leclerc et al. (2023) recently showed how combining a reaction-diffusion model and image data can improve plant-pathogen phenotyping. While such approaches are common in medical sciences (Mang et al., 2020), for instance in the study of tumor growth (Hogea et al., 2008), it remained unused in plant pathology. They were able to describe the overall spread of the visible lesion but pointed out a significant leaf deformation that was not taken into account. Besides improving the description of the interaction, considering plant growth and shape may be of great interest to study plant-pathogen interactions, for instance to identify new traits related to pathogen aggressiveness or plant resistance to disease.

Several methods have been used in addressing the plant growth and shape (Marconi and Wabnik, 2021), among which used L-systems (Prusinkiewicz et al., 2018), dynamics in mesh (Jeong et al., 2013; Hong et al., 2005), level set (Pineda and Gwun, 2017a,b), and fourth-order PDE (Chaudhry et al., 2019). While some of these approaches can produce impressive results, for instance in virtual worlds, they require important computing resources or be difficult to test against empirical data. Following the work of Leclerc et al. (2023), the main methodological question is to reconstruct continuous shape deformations from image sequences without a strong knowledge regarding the involved biological processes. In this context, the use of level set presents many advantages to address this question.

The level set method, developed by Osher and Sethian in the 1980s is one numerical technique used to track and represent the evolution of interfaces or boundaries. It was initially introduced in the paper of Osher and Sethian (1988) which demonstrated the method’s application to problems involving propagating fronts, which could change topology and merge or split naturally. Since then it has attracted studies in various applications from fluid dynamics (Sussman et al., 1994) and computer vision (Malladi et al., 1995) to medical imaging (Hogea et al., 2006). The level set method used to estimate contours change in time from images is the active contour model, specifically designed for handling complex topologies and capturing boundaries (Wang et al., 2021). Unlike other approaches, this method has more advantages in terms of topological flexibility (Osher and Sethian (1988)), implicit representation (Sethian, 1999), numerical stability and robustness (Osher and Fedkiw, 2001), ease of implementation (Chan and Vese, 2001), and a unified framework for different problems (Malladi et al., 1995), making it a powerful and versatile tool in various applications, particularly in image processing and medical imaging.

In this study, we use the level set approach to monitor both leaf deformation and lesion growth. While such approach has already been used in biomedical sciences (Hogea et al. (2006)) it remains rarely considered in plant pathology. We consider the spread of *Peyronellaea pinodes* (formerly *Mycosphaerella pinodes* and *Didymella pinodes*) on pea stipules (i.e. unconventional leaves in so-called semi-leaflet pea cultivars) as an example pathosystem. To assess how level set method can improve the phenotyping of plant-pathogen interactions we use the image sequences data used by Leclerc et al. (2023).The synergy between image processing and level set methods results in a powerful tool for tracking lesion growth in plants. This combination leverages the strengths of both approaches to provide accurate, robust, and detailed analysis of leaf and lesion evolution. The combined approach provides clear visual feedback on lesion boundaries and their evolution over time, aiding in better understanding of plant health status. After a brief presentation of the image sequences data acquired during the longitudinal monitoring of lesions on inoculated leafs and extraction of leaf and lesion contours from the pixel-segmented and probability images, we introduce the model proposed by Bertalmio et al. (2000) and Vemuri et al. (2003) derived from the level set formulation. This approach is then used to track both leaf deformation and lesion growth from the images. We show that this approach is efficient in capturing these phenomena from image data and study plantpathogen interactions. We finish by discussing further developments to improve the processing of images acquired by plant pathologists and to reduce further discrepancies observed in the solution of the method.

## 2. Materials and Methods

In this study we considered the data used by Leclerc et al. (2023). The dataset consists of images of pea stipules inoculated with an aggressive isolate of *Peyronellaea pinodes*, a fungal pathogen that belongs to the Ascochyta blight of pea disease complex that causes substantial yield losses (Bretag et al., 2006; Dutt et al., 2020). The data includes two pea cultivars with contrasted level of quantitative resistance: Solara, susceptible to infection by *P. pinodes*, and James, that exhibits partial or quantitative resistance in controlled conditions (Onfroy et al., 2007). For each cultivar, 16 inoculated leaves were monitored daily with visible imagery between 3 and 7 days after inoculation. For each sequence, images were registered (i.e. aligned to each other) assuming rigid transformations with the first image (3 days after inoculation) as the reference. Afterwards, images were segmented by classifying pixels in either healthy, symptomatic or background states with a validated supervised method (Jumel et al., 2022). RGB, segmented and probability images, giving the probabilities of each pixel to be in each state, are available in the open dataverse (Leclerc et al., 2022).

### 2.1. Extraction of leaf and lesion contours

As we were interested in monitoring leaf deformation and lesion spread, we used these images to extract both leaf and lesion contours. In details, leaf contours were easily obtained from the segmented images with merging healthy and symptomatic pixels. On the other hand, for the lesions, boundaries are less sharp as appearing in the pixel-segmented images and probability images, making them more difficult to detect. Hence, we applied the marching squares algorithm (Lorensen and Cline, 1987) on probability images, giving the predicted probability of each pixel to be symptomatic to extract the contour of the lesion, assuming the visual lesion contour is given by probability 0.5 (S1 Appendix).

Leaf and lesion contours are illustrated in Figure 1 plotted in RGB image sequences. All contours were used to produce binary masks of both leaves and lesions (S2 Appendix).

**Figure 1:**
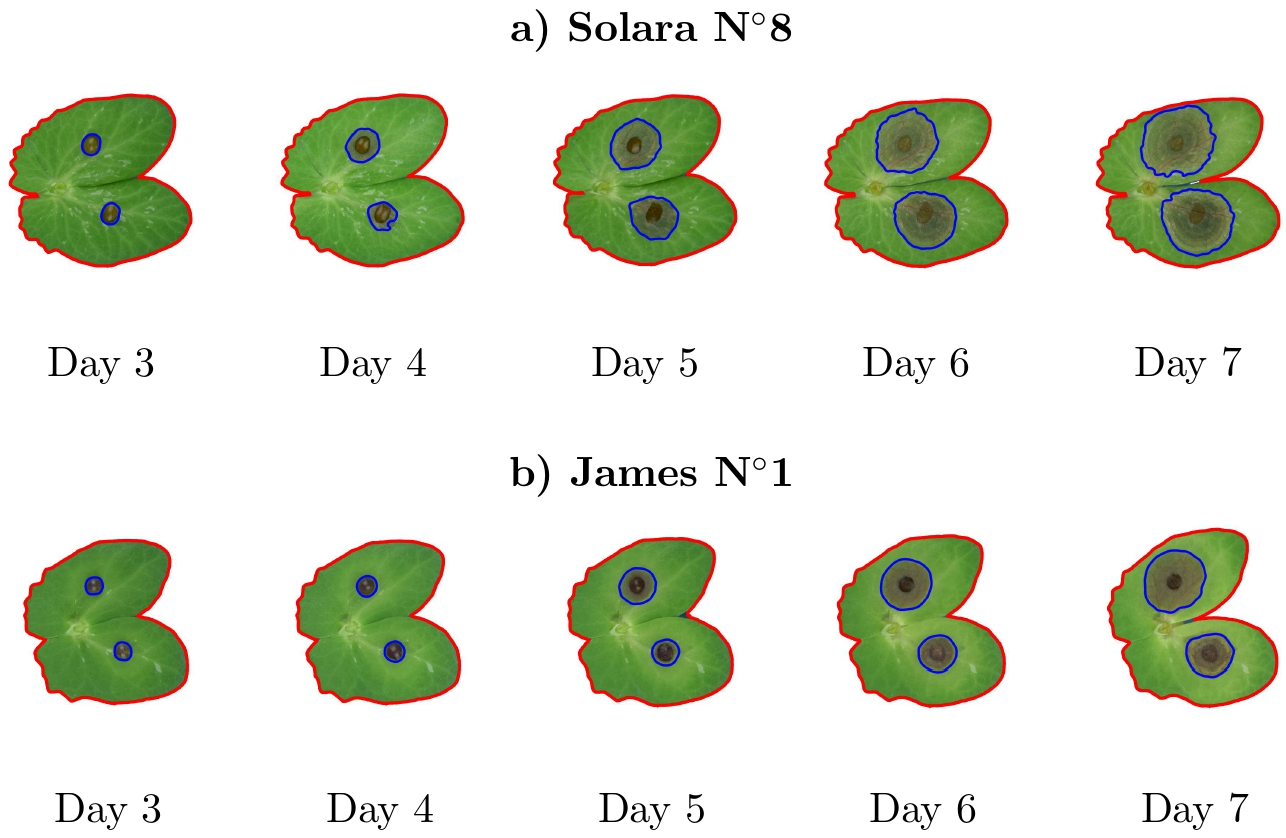
Leaf contours (red) and lesion contours (blue) on registered images of (a) Solara and (b) James cultivars.

### 2.2. Level set method for tracking contour evolution in time

The level set method allows to track and analyze the evolution of contour over time in a sequence of images. Various studies have utilized the level set method for contour evolution in different fields especially in medical imaging and computer vision. The level set method stands out in tracking contour evolution in medical imaging due to its adaptability to intensity inhomogeneity and robust segmentation results (Feng and Bing, 2022; Wang et al., 2021). It effectively handles complex topologies and boundary capture, making it particularly suitable for medical images with intensity variations (Yunyun et al., 2020).

As illustrated in Figure 2, we denote by *ϕ* the level set function such that the contour is the zero level set of *ϕ* defined as

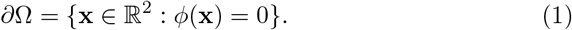

Moreover, the image surface is Ω = {x) ∈ ℝ^2^ : *ϕ* (x) < 0} and its complement is Ω^*c*^ = {x ∈ ℝ^2^ : *ϕ* (x) > 0} (Sethian, 1999; Osher and Fedkiw, 2002).

The contour evolution is given by the following advection equation, for *t* ∈ [0, *T*] and x ∈ Ω:

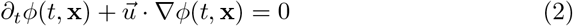

where *ϕ* is the level set function with the contour defined in equation (1), ∂_*t*_*ϕ* is the change in the level set function, ∇*ϕ* is the gradient of *ϕ* representing the rate of change of *ϕ* in space, and 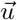 is the velocity field that dictates how the level set function *ϕ* changes over time. It is important to note that the contour evolution considers the motion of the contour in the normal direction, that is, the exterior normal vector to the contour represented by the level set function *ϕ* is given by 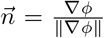 where ∥∇*ϕ*∥ is the magnitude or norm of the gradient vector ∇*ϕ*.

**Figure 2:**
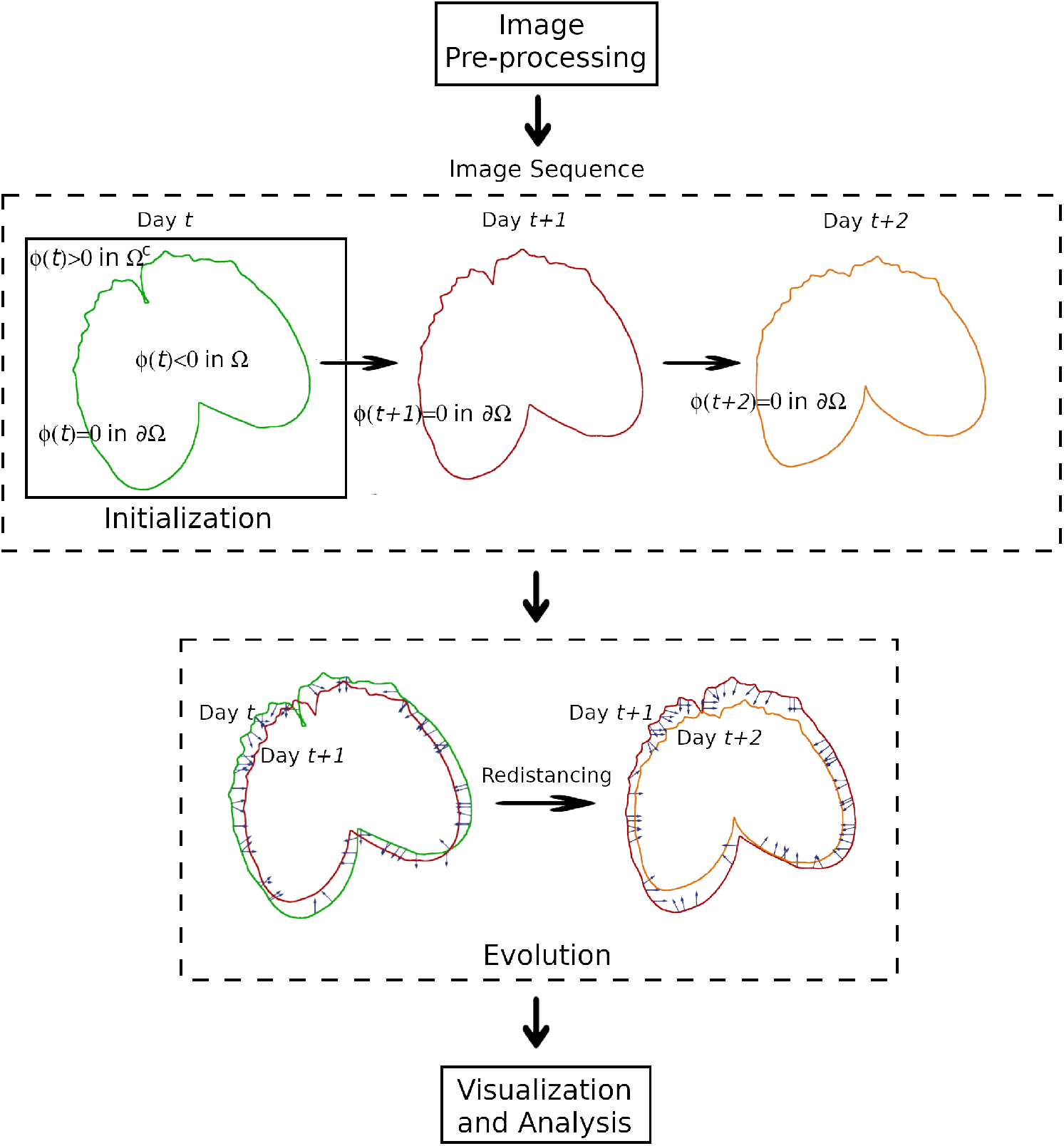
Schematic representation of level set method in tracking contour evolution between successive days.

The velocity field is a crucial component in the simulation, guiding how the size of the image evolves. We choose to compute the velocity field using the signed distance function between two successive times, an approach based on the mathematical models proposed by Bertalmio et al. (2000); Vemuri et al. (2003):

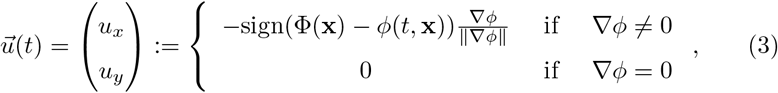

that is, the function *ϕ* (*t*, x) is morphing to the function Φ (x) following the normal direction.

The gradient of the level set function *ϕ* is approximated by a second-order centered finite difference scheme. If we denote *ϕ*_*i,j*_ the approximation of *ϕ* at **x** = (*x*_*i*_, *y*_*j*_), and Δ*x* and Δ*y* the spatial step sizes in the *x* and *y* directions, respectively, we write

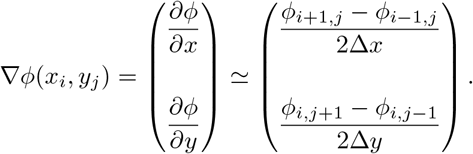

Once the velocity field 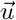 is computed, the advection equation 2 is solved. The upwind differencing scheme for the advection equation is given by, for 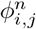 the approximation of *ϕ* at time *t*_*n*_ and x = (*x*_*i*_, *y*_*j*_):

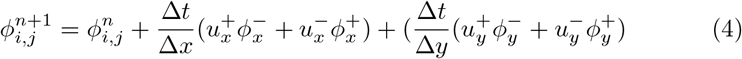

with

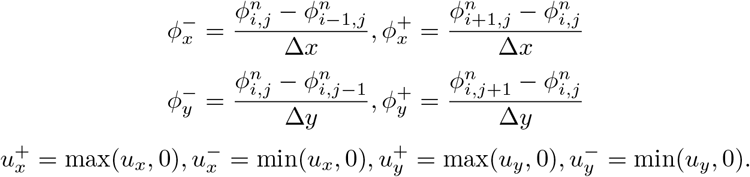

In equation 4, Δ*t* is the time step size,, 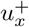 and 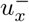 are the positive and negative parts of the velocity component in the *x* direction, 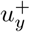 and 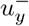 are the positive and negative parts of the velocity component in the *y* direction, 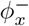 and 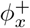 are the values of *ϕ* at the neighboring grid points in the *x* direction, and 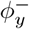 and 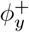 are the values of *ϕ* at the neighboring grid points in the *y* direction. Finally, the time step Δ *t* is chosen to satisfy the CFL stability criterion:

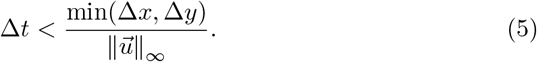

where 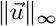 is the maximum norm of the velocity field (Osher a nd Sethian, 1988).

Due to various operations performed in the simulation, as time progresses, the signed distance property may be lost, and the function may no longer represent a signed distance function to the interface. Hence, redistancing is performed periodically to restore this property and maintain accuracy, ensuring that it closely matches the true boundaries (Osher and Fedkiw, 2002).

### 2.3. Application to leaf deformation and lesion growth

Here we used the level set method presented above in Figure 2 to track the evolution of leaf and lesion contours through time using sequence of images (Figure 1). For each inoculated leaf, we started from the extracted leaf and lesion contours. We used the level set method to track contour changes between day 3 and day 7; hence, the initial image is at Day 3. For each image sequence there were five images. The level set method is implemented repeatedly from Day 3 to Day 4, Day 4 to Day 5, and so on until Day 6 to Day 7. In effect, the implementation of the method started on the first day the lesion was detected (S1, Appendix). Once the contour evolution process is complete, we analyze the tracked contours in each frame to extract quantitative information about the image’s behavior over time. Outcomes in the implementation are presented by visualization and metric measures in comparing the image data and the level solution as well as comparing between different cultivars.

The adequacy of the model to the contours data were assessed visually using Paraview (Ahrens et al., 2005), an open source visualization software. Moreover, the quality of the model was quantified using two metrics: the Jaccard Similarity Index and the Relative Error. The Jaccard Similarity Index is utilized to measure the similarity between sets by comparing the intersection over the union of the segmented regions (Kamiura, 2003). It ranges from 0 (no intersection) and 1 (perfect matching), and stands out in image analysis compared to other distance metrics due to its effectiveness in calculating similarity between images based on positional feature vector comparisons, indirectly considering shape, position, orientation, and other features (Gonzalez-Huitron et al., 2003). On the other hand, the relative error was computed using the *L*^1^-norm given by

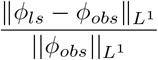

where *ϕ*_*ls*_, *ϕ*_*obs*_ represents the level set solution and the observed image, respectively, and 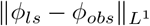 is the *L*^1^ -norm of the difference between the level set solution and the data image. Differences between days and cultivars on these two metrics were tested statistically using Analysis of Variance (ANOVA). Image processing and the level set method implementation were performed using the open programming language Python.

## 3. Results

### 3.1. Application to Leaf Deformation

As shown in Figure 3 and Figure 4 for two sample image sequence of leaves, in all cases a less noticeable difference between the observed images of the leaf and level set solution can be observed since black contours (level set solution) are overlapping the gray contours (observed images) in almost all parts of the contour. However, with respect to days of observation, more noticeable difference can be observed on Day 7 compared to the previous days. We observed similar results for all the set of image sequences (S3, Appendix).

**Figure 3:**
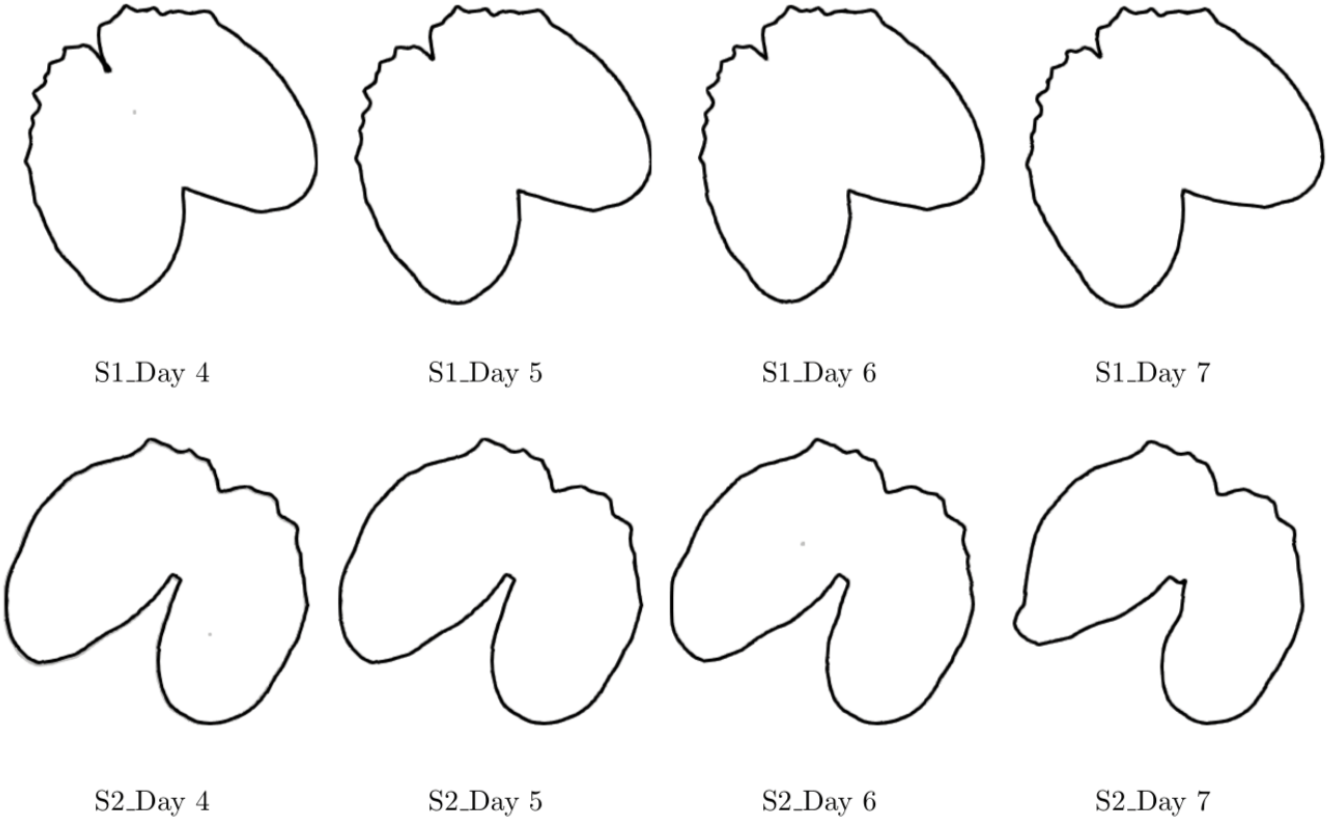
Two samples of image sequences, S1 and S2 of Solara cultivar comparing observed images of leaf (gray contour) and level set solution (black contour).

**Figure 4:**
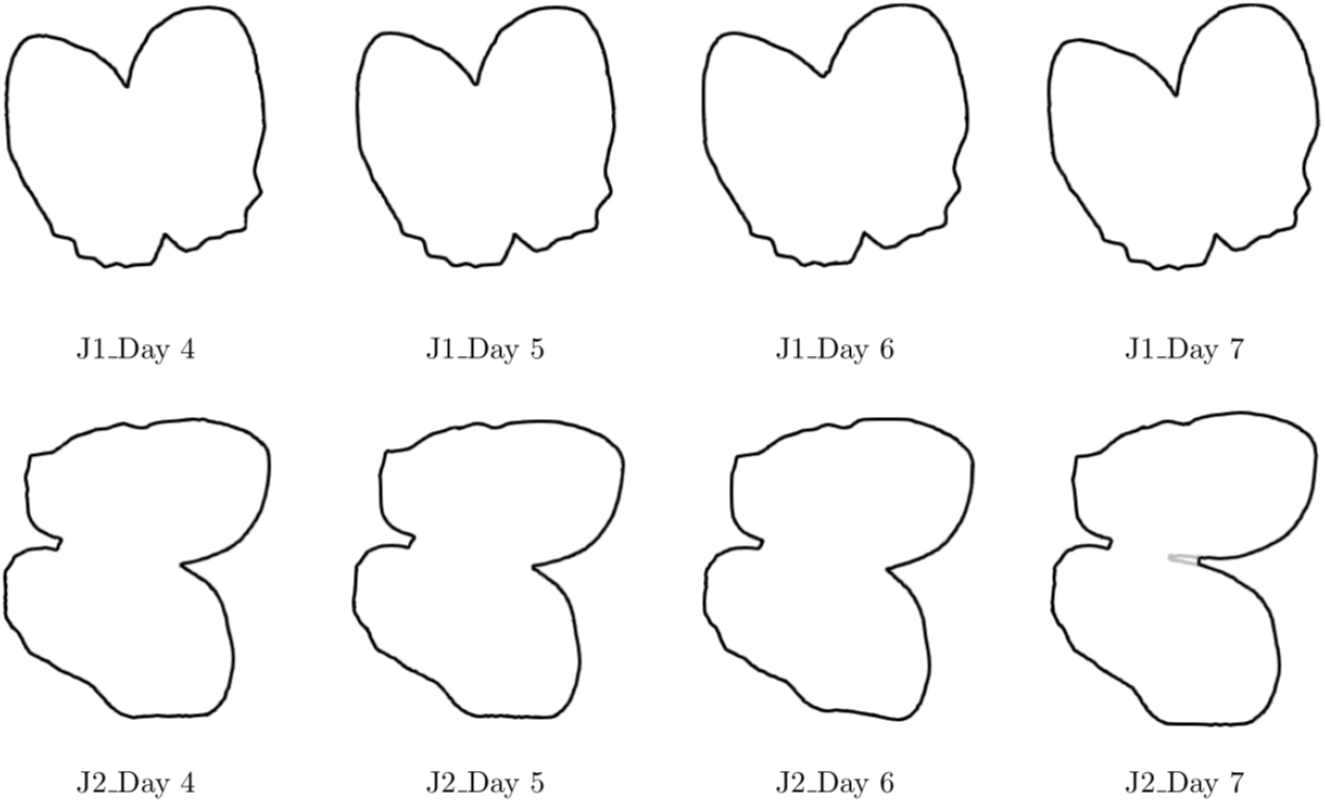
Two samples of image sequences, J1 and J2 of James cultivar comparing observed images of leaf (gray contour) and level set solution (black contour).

Overall, the Jaccard index shown pointed out a good similarity between the model and the observed leaf contours (S5.1, Appendix). It ranged from 0.869 (Day 7 of Solara N°2) to 0.999 (Day 4 of Solara N°9)for Solara cultivar while it was between 0.938 (Day 7 of James N°14) and 0.99 (Day 4 of James N°14)for James, suggesting a higher variability for Solara cultivar than for James. The visualization of the Jaccard index against the date in Figure 5 showed a slow decrease between Day 4 and Day 7 for both cultivars that was found to be statistically significant (p-value < 0.05). Furthermore, the statistical analysis also highlighted a significant interaction between date and cultivar (p-value < 0.05), only explained by a lower index for Solara at Day 7.

**Figure 5:**
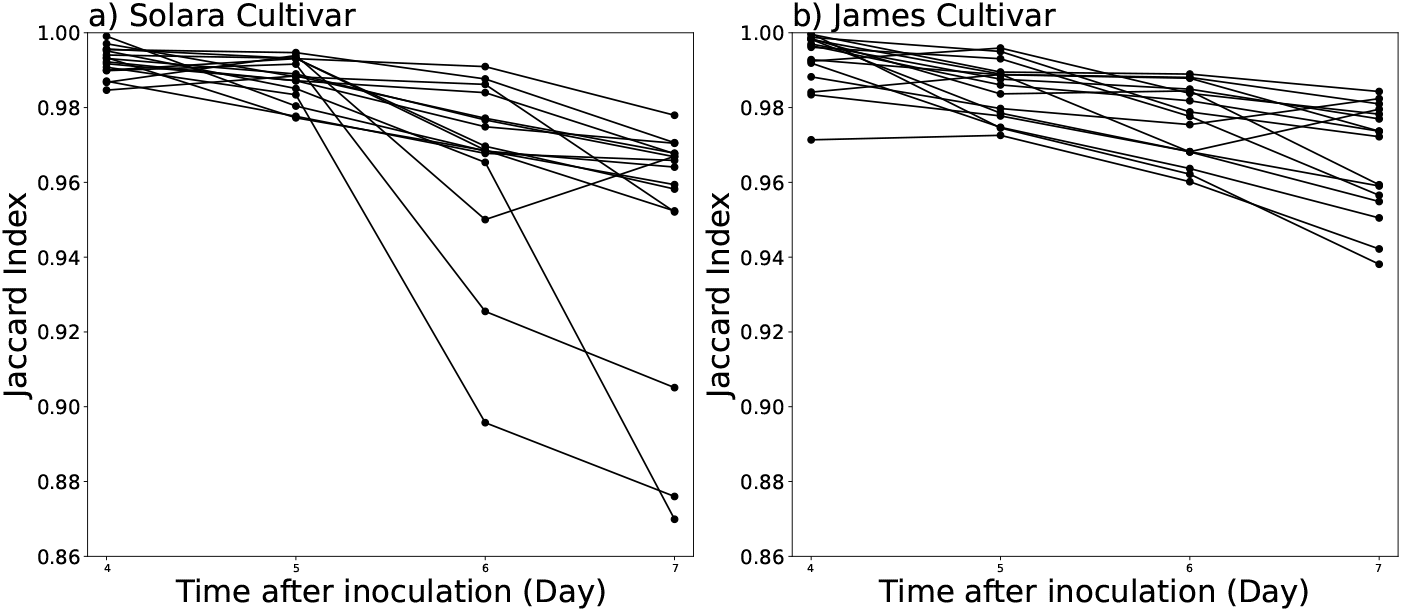
Change in the Jaccard similarity index in leaf deformation with time for Solara and James cultivars.

As depicted in Figure 6, James cultivar had a higher mean relative error value (0.07) compared to Solara cultivar equivalent to 0.03. Moreover, outliers are present in both cultivars with two outliers for Solara cultivar coming from Solara N°2 (0.07) and Solara N°12 (0.12) images and three outliers for James cultivar coming from James N°1 (0.132), James N°8 (0.05), and James N°13 (0.134) images (S6.2, Appendix). The analysis of variance only suggested a significant effect of the date (p-value < 0.05) that was explained by significant higher errors at Days 5 and 6. However neither the cultivar nor the interaction between the date and cultivar were found significant (p-value > 0.05). Relative error values for all 32 image sequences across days are available in the supplementary material (S6.1, Appendix).

**Figure 6:**
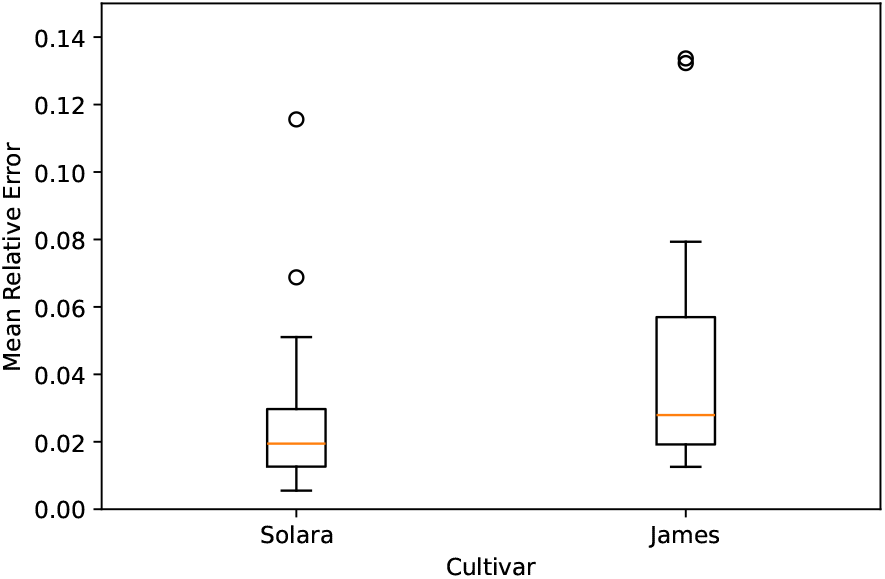
Boxplot of relative errors computed comparing the level set simulations and observed leaves of Solara and James cultivars.

### 3.2. Application to Lesion Growth

Similarly to leaf deformation, and as illustrated in Figure 7 and Figure 8 for the four sample lesions, the level set solution reproduced the contours of the lesions with a good accuracy. This is consistent with all of the 32 images sequences (S4, Appendix).

**Figure 7:**
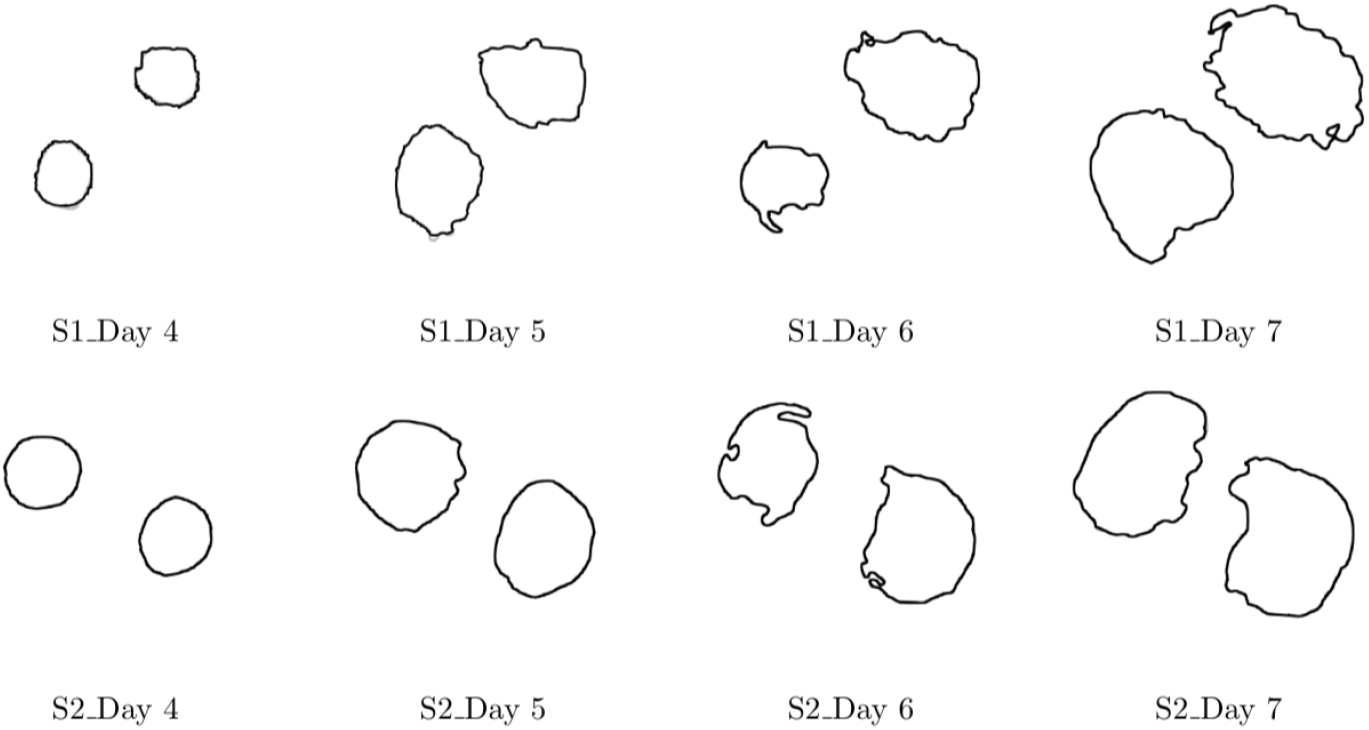
Comparison between observed images of lesion (gray contour) and level set solution (black contour) on Solara cultivar

**Figure 8:**
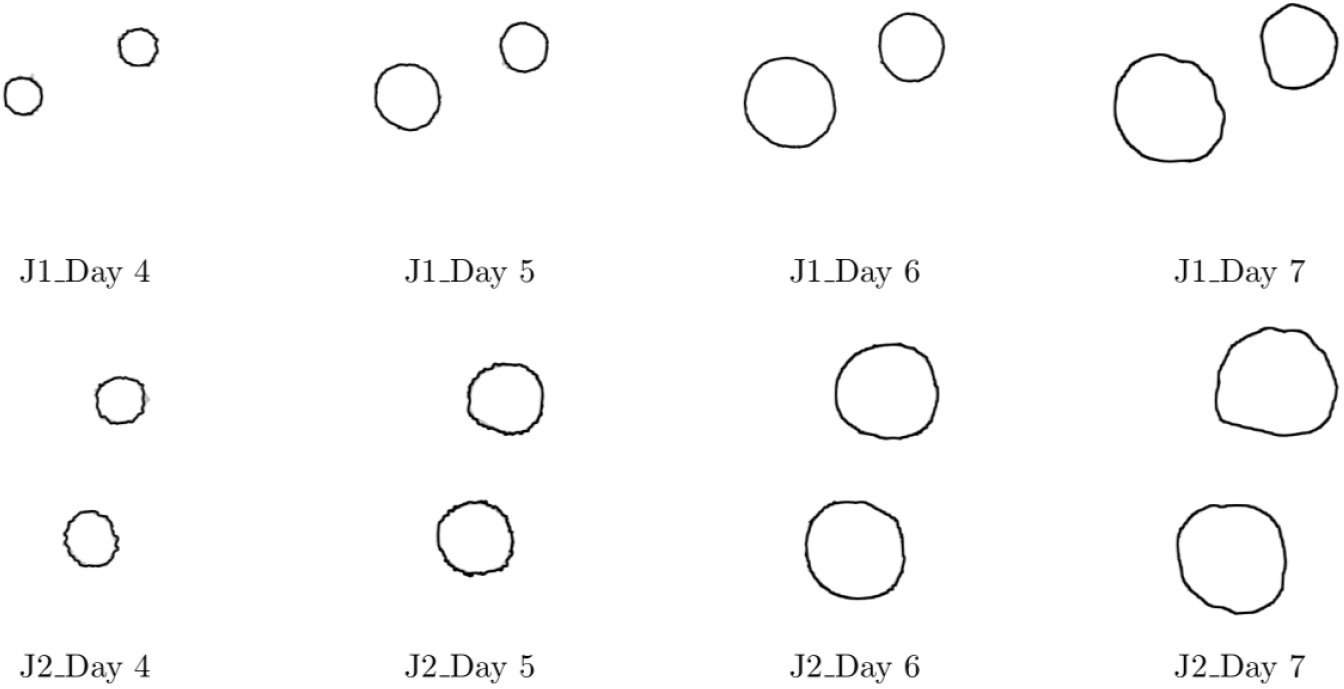
Comparison between observed images of lesion (gray contour) and level set solution (black contour) on James cultivar

The Jaccard index values as shown in Figure 9 were in agreement with the visual assessment of model adequacy. It ranged from 0.81 (Day 6 of Solara N°1) to 0.99 (Day 6 of Solara N°5) for Solara and between 0.96 (Day 3 of James N°1) to 0.99 (Day 7 of James N° 10) for James (S5.2, Appendix). The analysis of variance pointed out non significant effects of the cultivar and the date (p-values > 0.05). It only highlighted a significant interaction between date and cultivar (p-value < 0.05), only explained by a lower index for Solara at Day 6.

**Figure 9:**
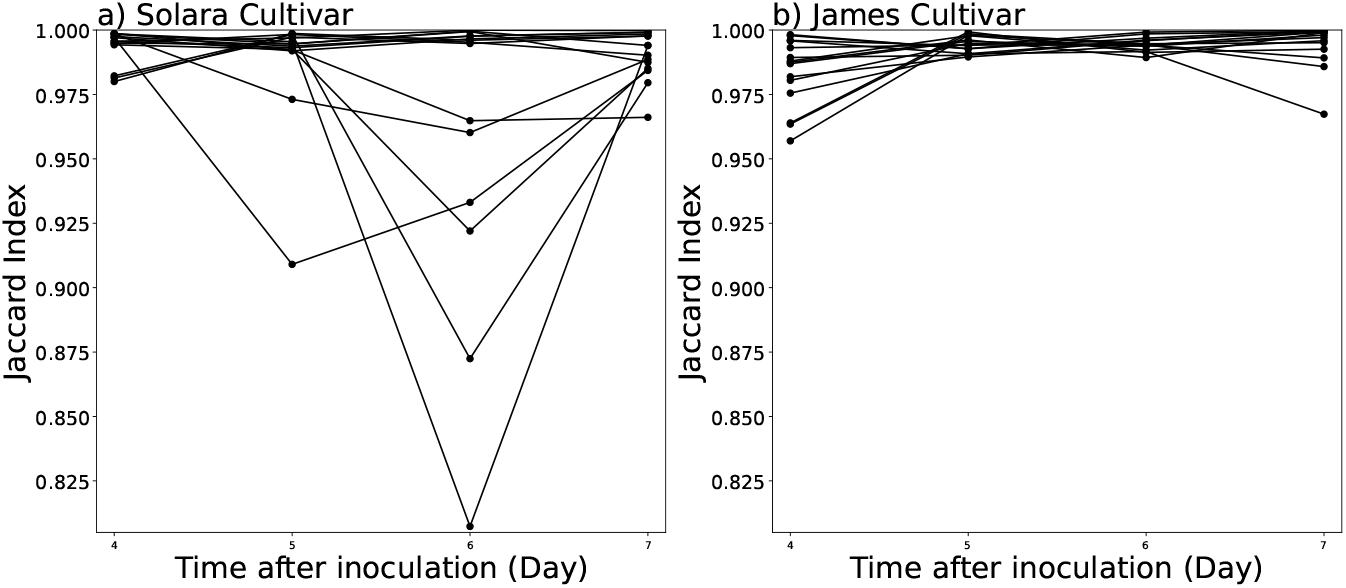
Change in the Jaccard similarity index for lesion growth with time of Solara and James cultivars

The relative error values as depicted in Figure 10 were also in agreement with a good performance of the model. The visualisation of the overall values showed James cultivar had a higher mean relative error (0.03) compared to Solara cultivar (0.01). Two outliers were observed in Solara cultivar coming from Solara N°9 (0.03) and Solara N°11 (0.09) images while one outlier is noticed for James cultivar coming from James N°5 (0.12) image (S6.4, Appendix). However the statistical test only exhibited significant effects of the date and its interaction with cultivar (p-values < 0.05), the suggested overall difference between cultivars being non significant (p-value > 0.05). Moreover, relative error values for all 32 image sequences across days are available in the supplementary material (S6.3, Appendix).

**Figure 10:**
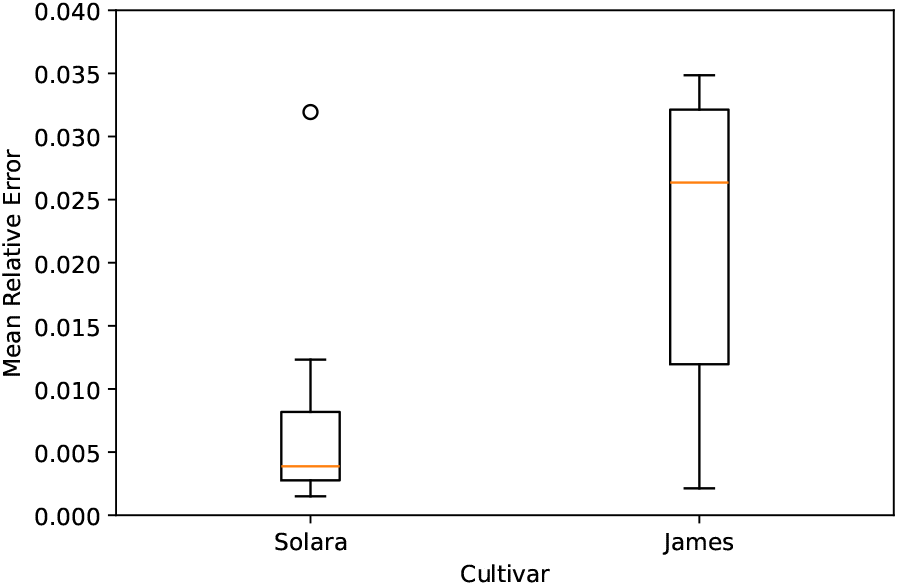
Boxplot of relative errors computed comparing the level set simulations and observed lesions of Solara and James cultivars.

## 4. Discussion

Considering the spread of *P. pinodes* on pea stipules, the implementation of the level set method along with its numerical implementation (Osher and Fedkiw (2002); Bertalmio et al. (2000)) allowed us to track the evolution of the leaf deformation and lesion growth. This combination of image-based phenotyping and spatial mathematical modeling was able to provide a new approach in describing plant-pathogen interactions when pathogen induces growing lesions. Although this method has been used to model the dynamics of shapes in many fields such as medical imaging and diagnosis (e.g.Swierczynski et al. (2018); Mang et al. (2020)), this approach remains poorly considered in plant pathology. Level set approaches are interesting methods that researchers in plant pathology should consider in their toolbox for processing image-based phenotyping data. Here we utilized the image sequences data used by Leclerc et al. (2023) to extract leaf and lesion contours with its evolution described by the advection equation and the contour defined by the level set function. Using the PDE model and ensuring redistanciation periodically to maintain accuracy and numerical stability of the method, we were able to efficiently monitor both the leaf deformation and lesion spread in time.

In monitoring leaf deformation, minimal visual differences were observed between the image data and the level set solution, with more noticeable discrepancies on Day 7 compared to earlier days. The Jaccard Similarity Index indicated higher variability in Solara than in James cultivar. There was a significant decrease in the index from Day 4 to Day 7, highlighting Solara’s lower index on Day 7. James had a higher mean relative error than Solara, with outliers present in both cultivars. Significant errors were found on Days 5 and 6. However, neither cultivar nor the interaction between date and cultivar were statistically significant.

As lesion contours were difficult to extract from the images due to unclear interface, we used the marching squares algorithm (Lorensen and Cline, 1987) on slightly blurred probability images because it produced smoother lesions than others tested methods, e.g. level set segmentation on segmented images. Close alignment between the image data and the level set solution contours is visually evident and cultivar wise, James demonstrated higher accuracy in lesion growth tracking compared to Solara, with consistently higher Jaccard index values across days of observation. Both cultivars show variations in similarity between observed images and the level set solution over time, but Solara cultivar exhibits larger fluctuations compared to James cultivar. In addition, Solara cultivar has lower mean relative errors compared to James cultivar. However, the statistical test only showed significant effects for the date and its interaction with the cultivar, while the overall difference between cultivars was not significant.

The visual assessment of model adequacy and the two metrics used to evaluate the discrepancy between the model and the data pointed good overall performances. Although the numerical resolution of the advection equation can still be improved to reduce the minimal discrepancies found such as exploring higher order numerical schemes and smaller time steps, the model’s performance already provided a good convergence of the contours both for the leaf and lesion. Our approach allows a continuous reconstruction of shapes (i.e. leaf or lesion) from a few discrete observations (S1 and S2 Movies). This could be particularly interesting to provide realistic virtual spread of disease in existing virtual plant models (Garin et al., 2018). Moreover, in some cases the estimated vector field might have a biological interpretation (e.g. disease-induced deformation) and could provide original quantitative traits to describe plant-pathogen interactions. Even if it is based on a PDE equation, the approach used in this study does not require a mechanistic knowledge on the biological processes involved in the shape deformations and is easier to handle than other models that integrate more processes (Jeong et al., 2013; Hong et al., 2005). It could be interesting to compare different models to reconstruct plant leaf deformation and lesion spread in further works.

It is important to note that in this work, the level set was used to reconstruct shapes evolution and not to segment them. Of course, the modelled shapes evolution depends highly on the initial segmentation. For the leaves, the segmentation was easy as the interface between the stipules and the white background was obvious (for human and computer vision segmentation methods). However, the lesions exhibited less clear interfaces, especially for the last dates and as mentioned an algorithm aided in extracting the contours. In order to improve the description of the lesion it would be interesting to either improve lesion segmentation, knowing that it is even difficult to segment for an expert. Furthermore, one could introduce some mechanistic knowledge to provide a model for the vector field (Pineda and Gwun, 2017a), or to consider that the level set is the solution of a spatial mathematical model describing lesion spread. Finally, it would be worth coupling our approach with the reaction-diffusion model used by Leclerc et al. (2023) to fit the images. This would be particularly interesting to assess how taking into account a moving domain change the estimates of diffusion and reaction terms and the comparison between cultivars.

## Financial Support

This study is funded by Campus France and DOST-SEI under the PhilFrance-DOST Fellowship Program and the Plant Health and Environment Division of INRAE through the MODIM project. The funders had no role in study design, data collection and analysis, decision to publish, or preparation of the manuscript.

## Notes

https://entrepot.recherche.data.gouv.fr/dataset.xhtml?persistentId=doi:10.57745/MQXKCP

### Competing Interest Statement

The authors have declared no competing interest.

