## Supplementary Material for "Spatiotemporal Modeling of Host-Pathogen Interactions using Level Set Method"

### S1. Lesion contours

Contours shown in Figure 1 to Figure 8 were visualized using the raw images (RGB data) of the stipules for a clearer visualization. Since the level set method tracks moving interfaces, it is necessary that contours are present before its implementation. In the case of tracking the evolution of lesion contour, one image sequence of Solara cultivar (Solara N°15) has no detected lesion in Day 3 while another image sequence of James cultivar (James N °11) has no detected lesions in Day 3 and 4.

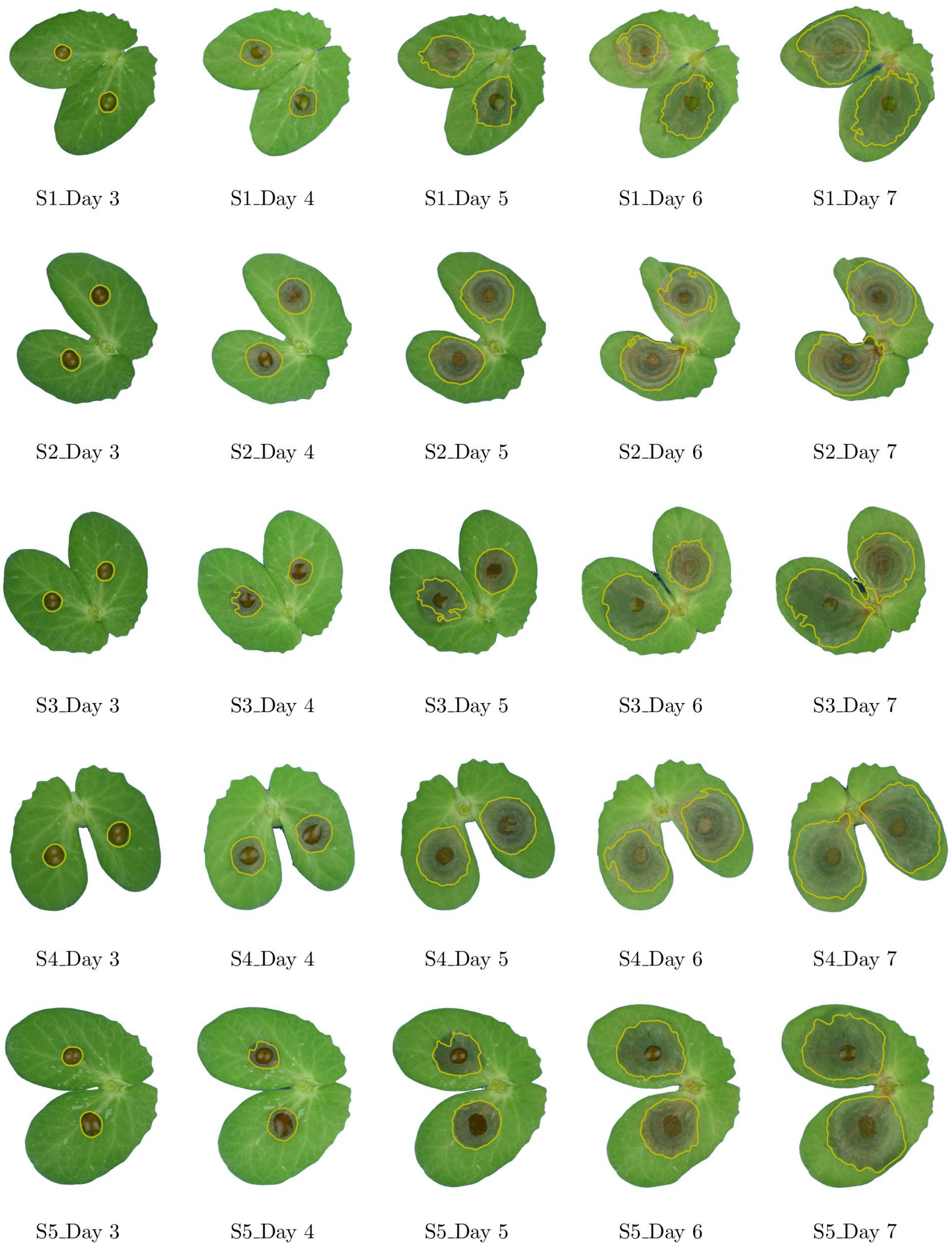


Figure 1: Lesion contours of *P. pinodes* on RGB images of Solara cultivar

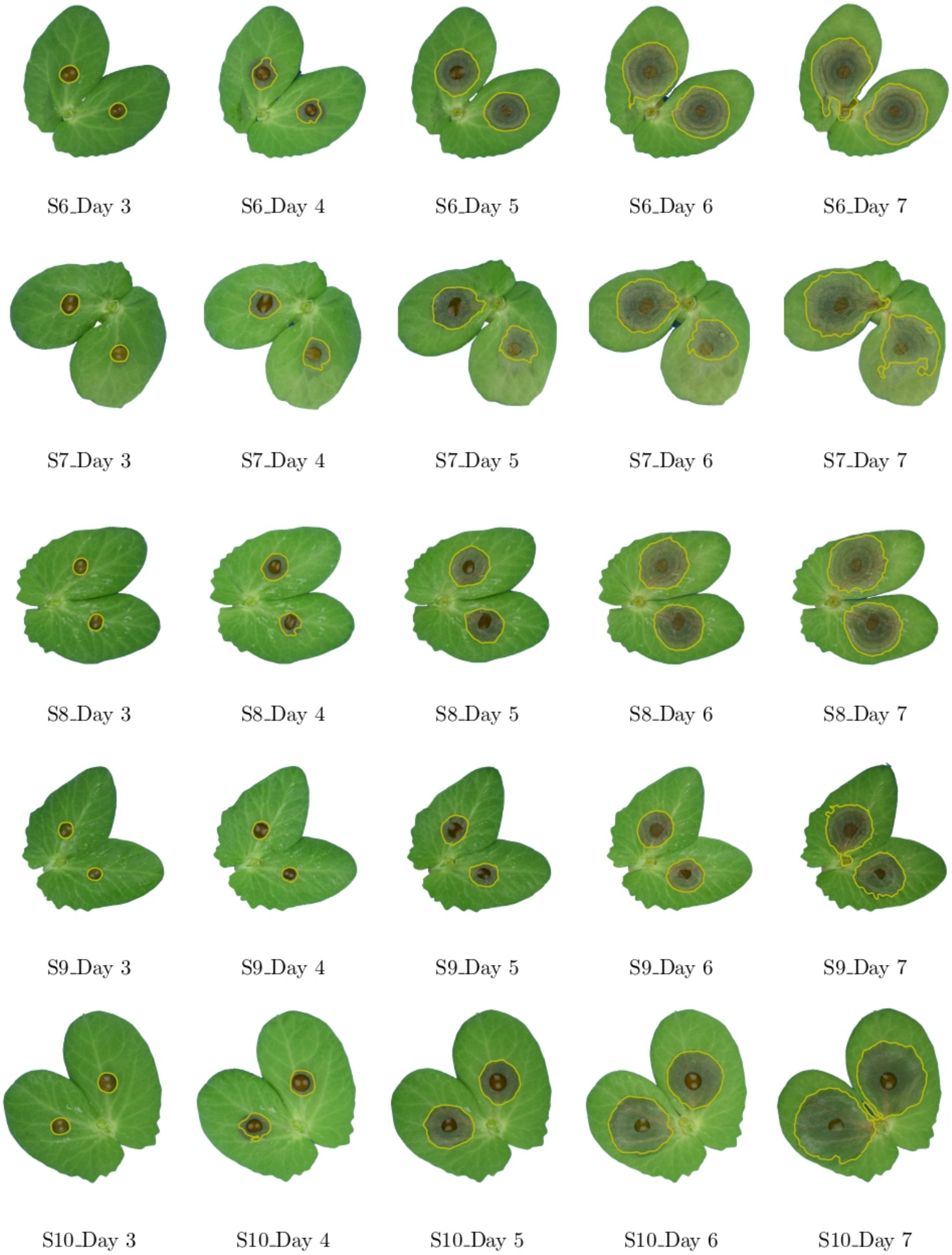


Figure 2: Lesion contours of *P. pinodes* on RGB images of Solara cultivar *(continued)*

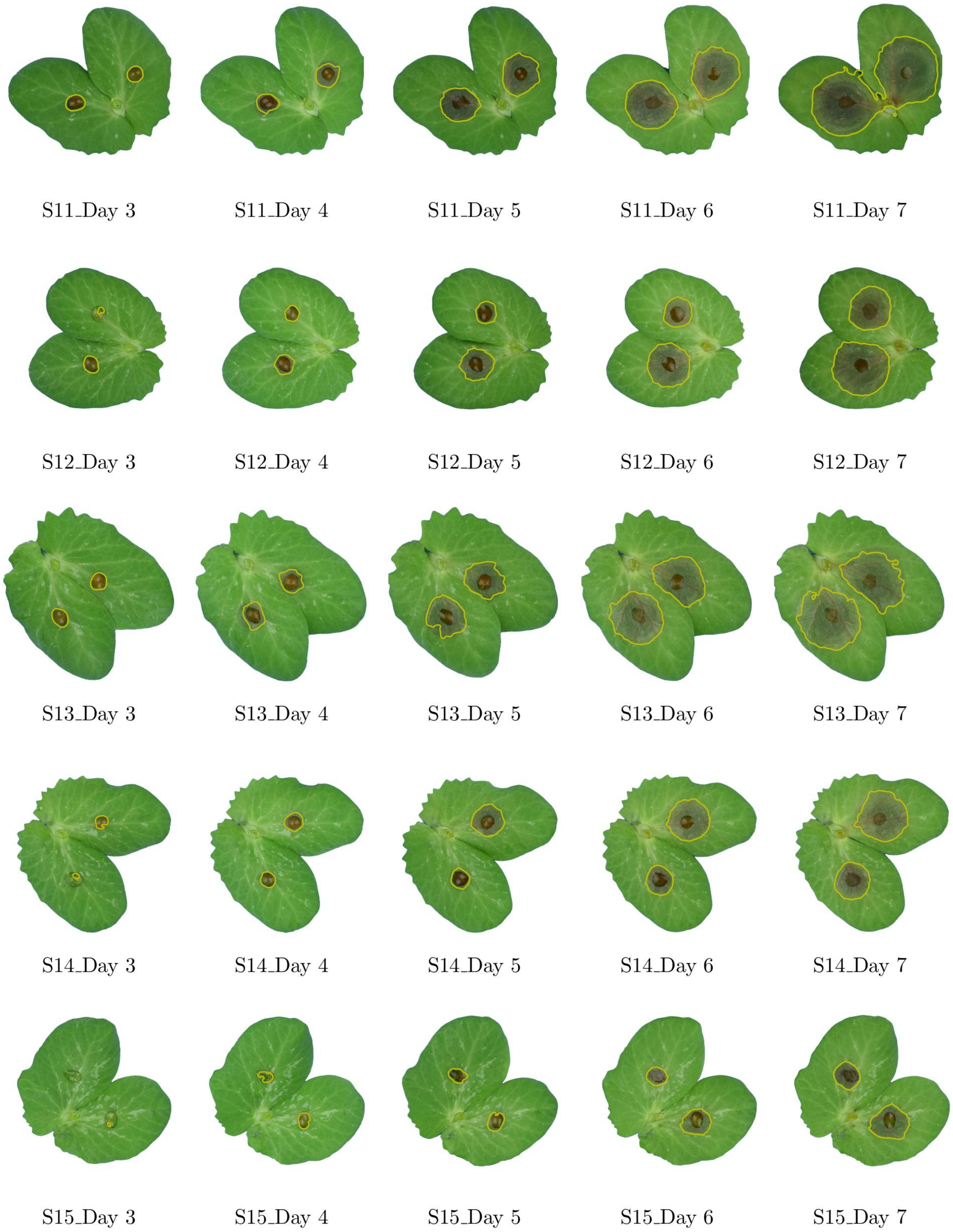


Figure 3: Lesion contours of *P. pinodes* on RGB images of Solara cultivar *(continued)*

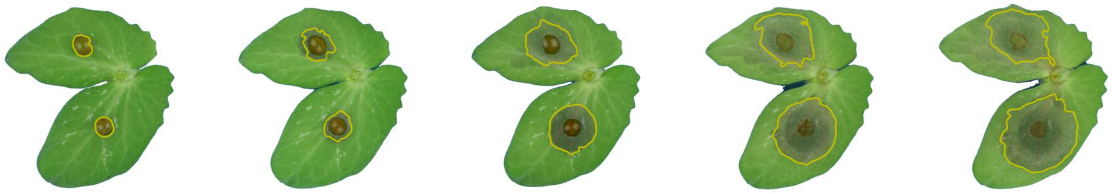

S16_Day 3 S16_Day 4 S16_Day 5 S16_Day 6 S16_Day 7

Figure 4: Lesion contours of *P. pinodes* on RGB images of Solara cultivar *(continued)*

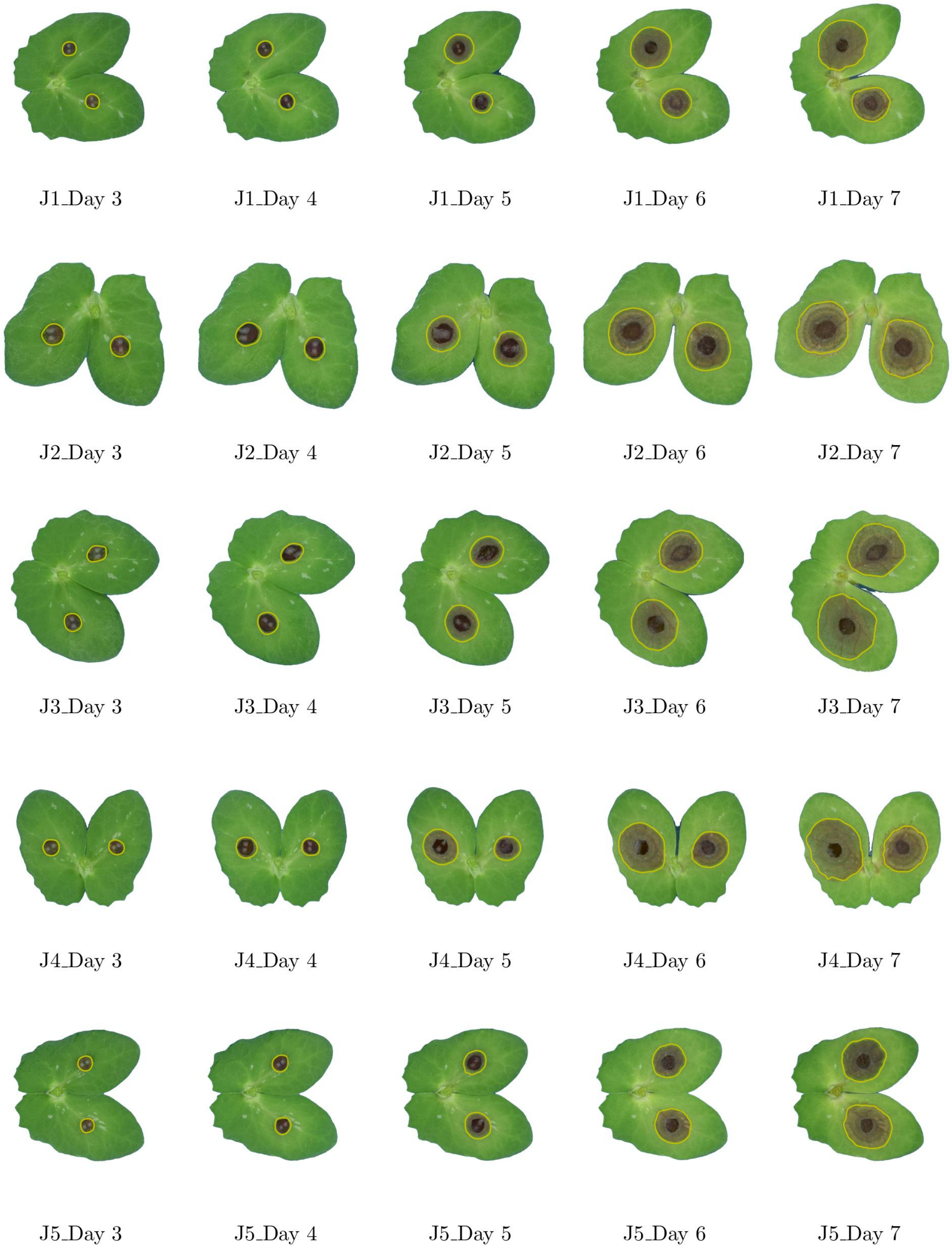


Figure 5: Lesion contours of *P. pinodes* on RGB images of James cultivar

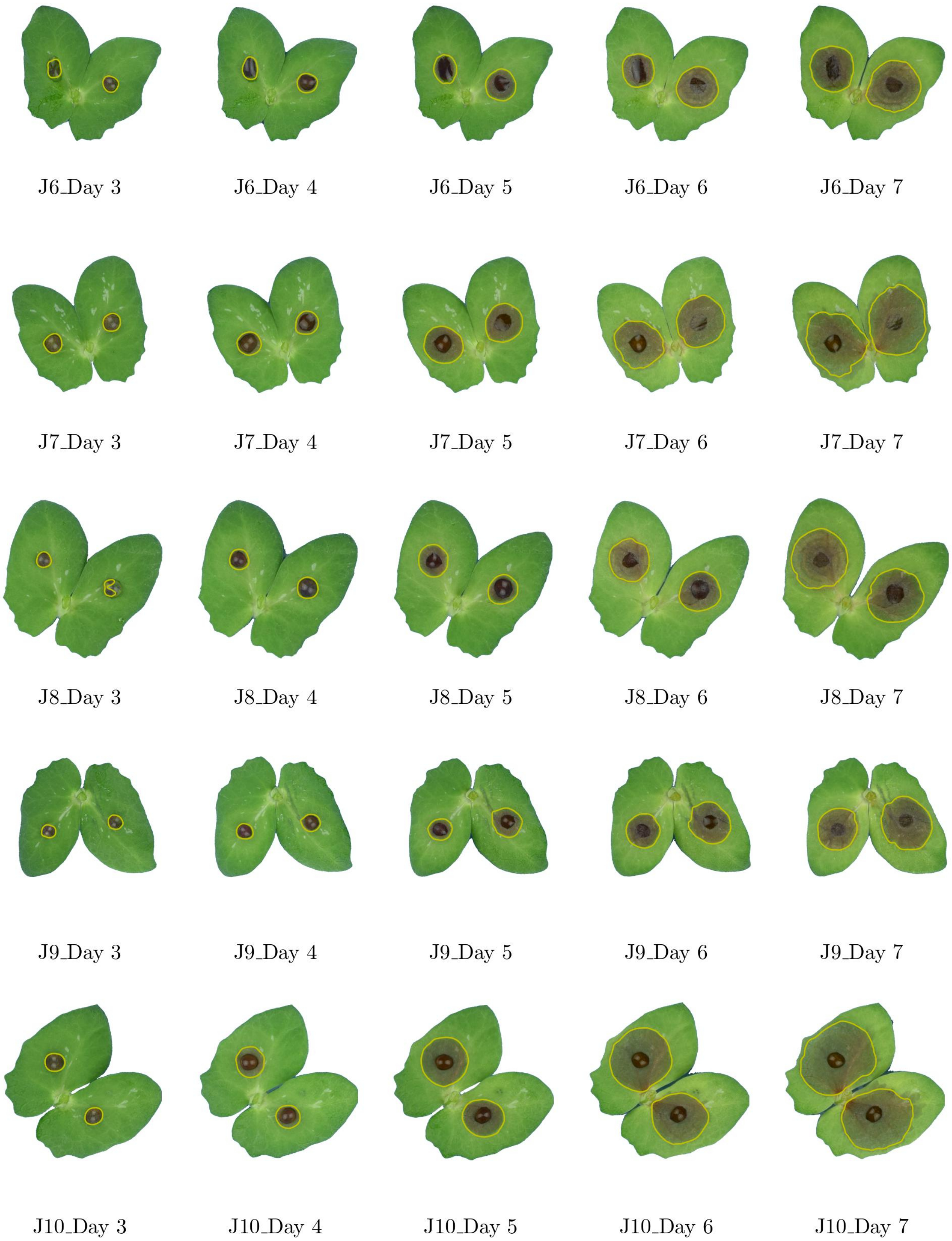


Figure 6: Lesion contours of *P. pinodes* on RGB images of James cultivar *(continued)*

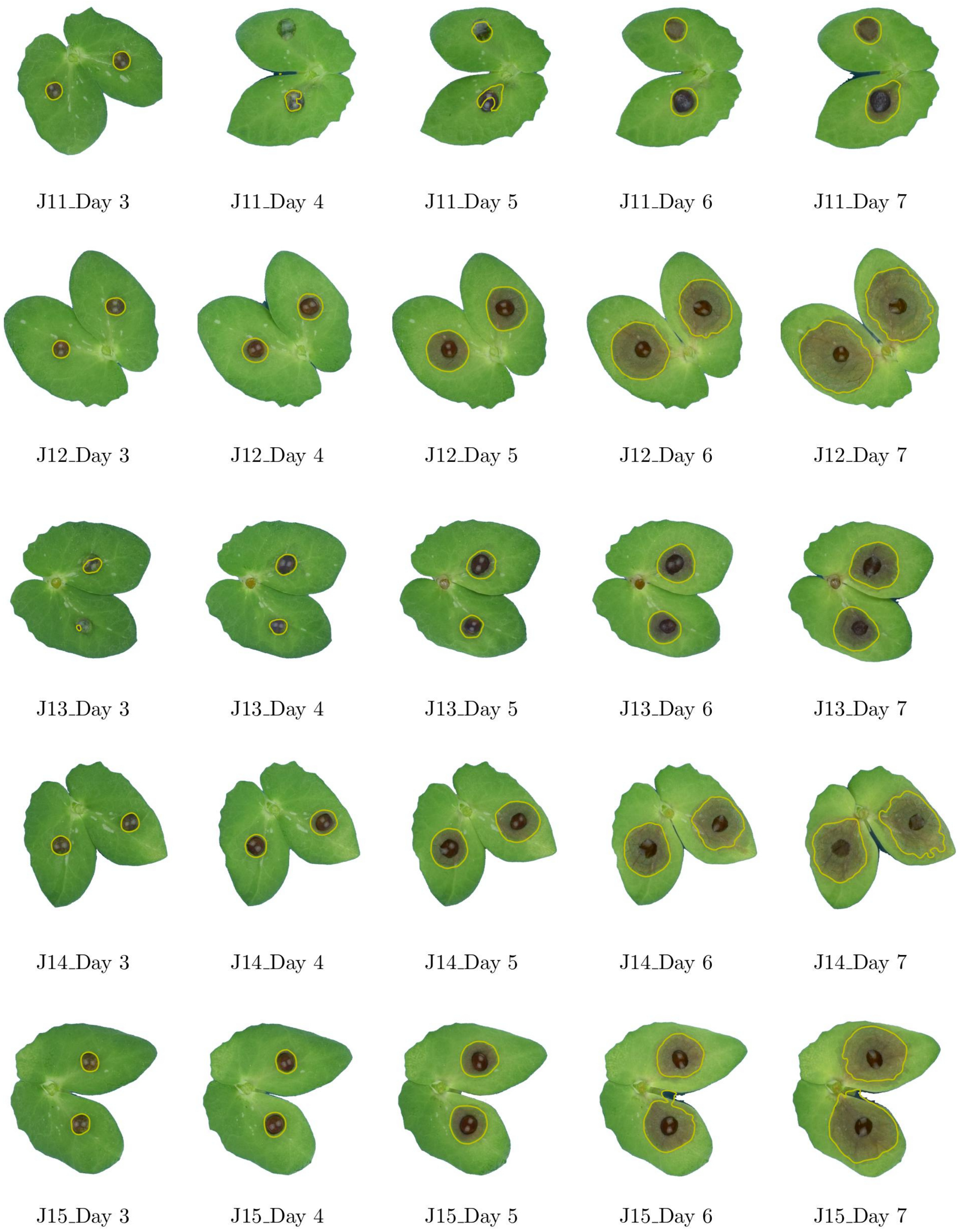


Figure 7: Lesion contours of *P. pinodes* on RGB images of James cultivar *(continued)*

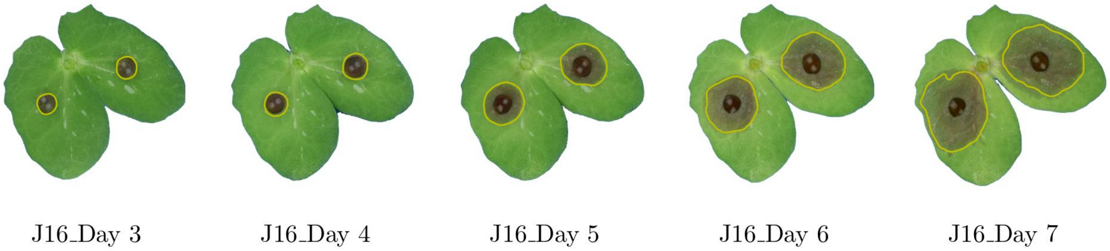


Figure 8: Lesion contours of *P. pinodes* on RGB images of James cultivar *(continued)*

**S2. Binary images from lesion contours**

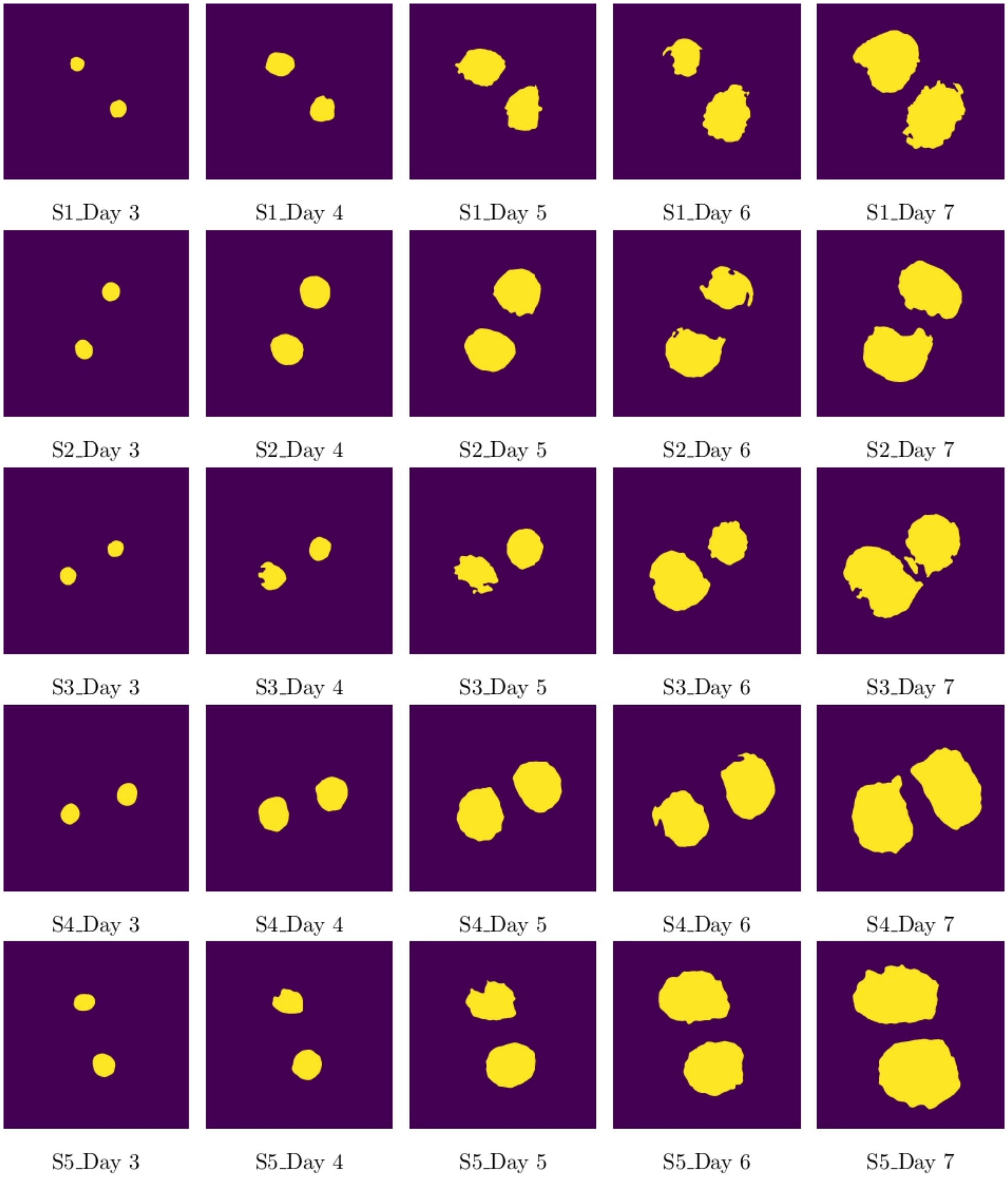


Figure 9: Binary images of lesion contours of *P. pinodes* on Solara cultivar

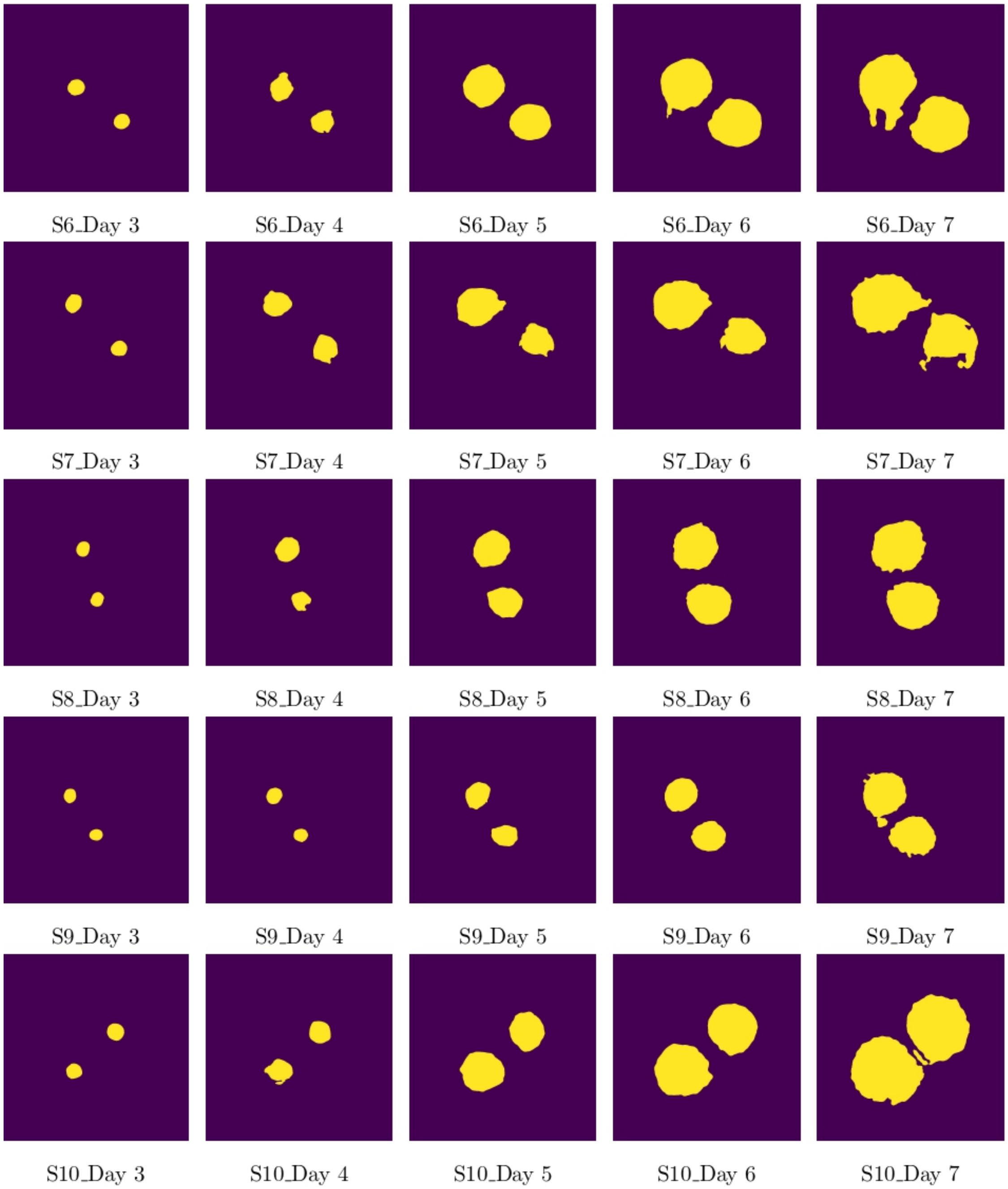


Figure 10: Binary images of lesion contours of *P. pinodes* on Solara cultivar *(continued)*

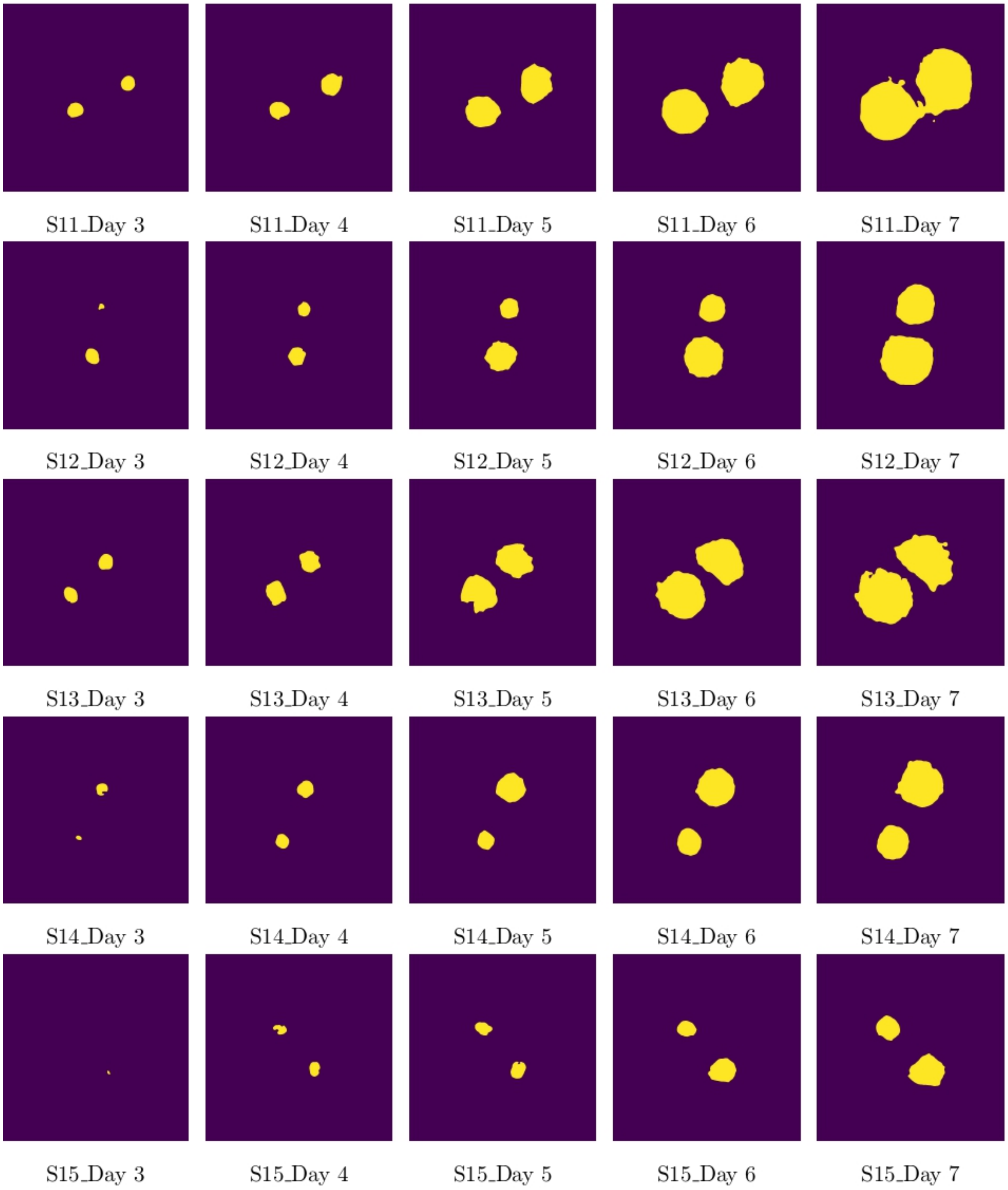


Figure 11: Binary images of lesion contours of *P. pinodes* on Solara cultivar *(continued)*

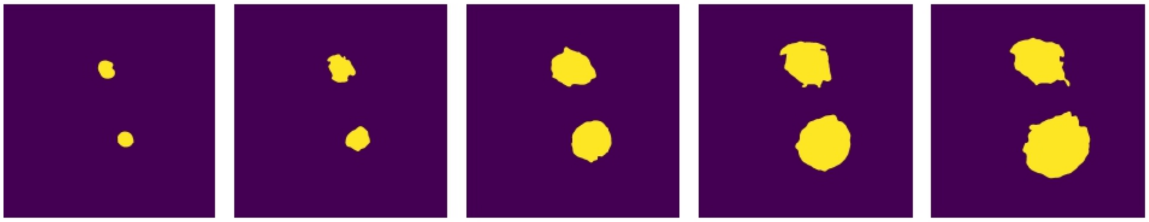

S16_Day 3 S16_Day 4 S16_Day 5 S16_Day 6 S16_Day 7

Figure 12: Binary images of lesion contours of *P. pinodes* on Solara cultivar *(continued)*

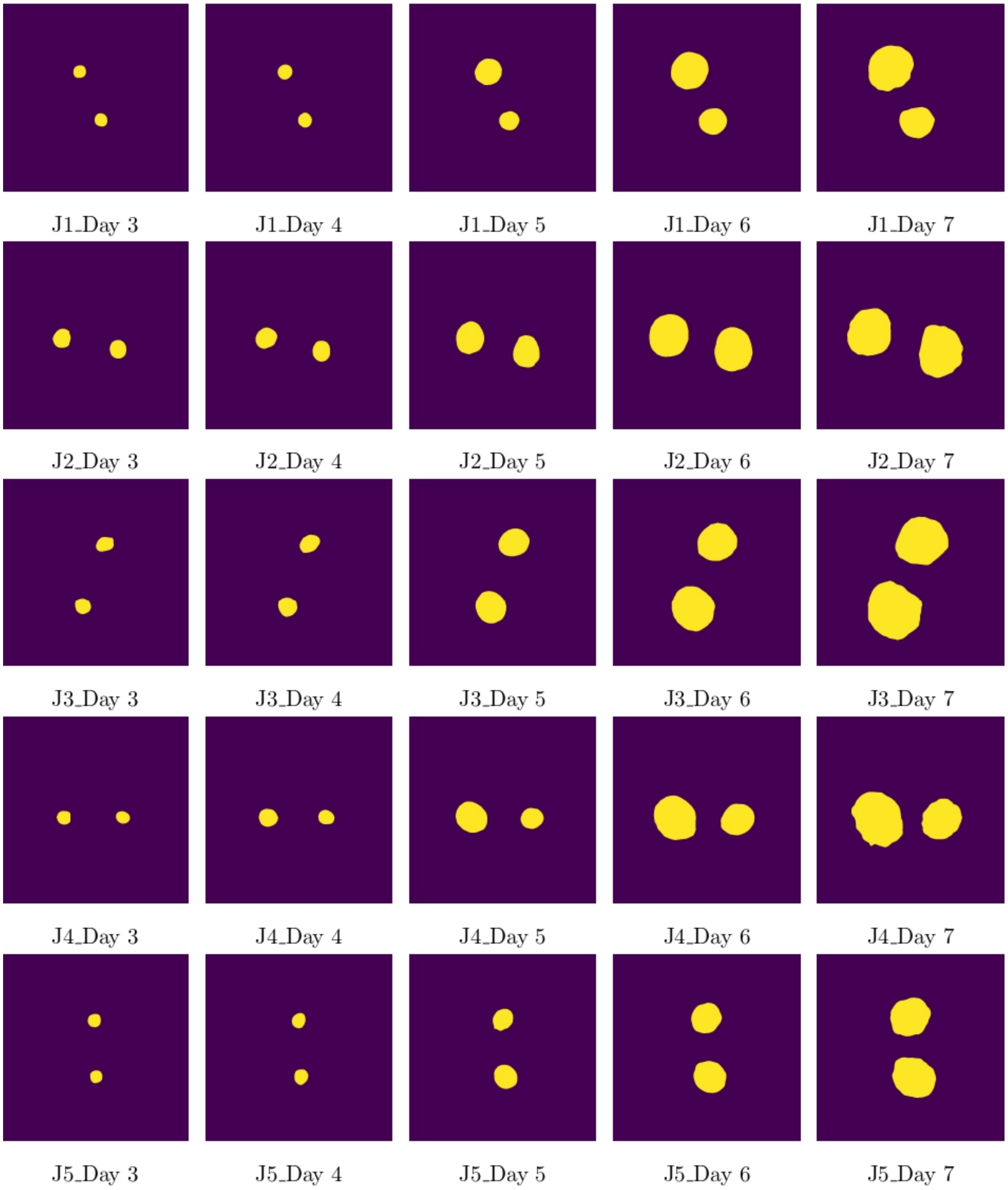


Figure 13: Binary images of lesion contours of *P. pinodes* on James cultivar

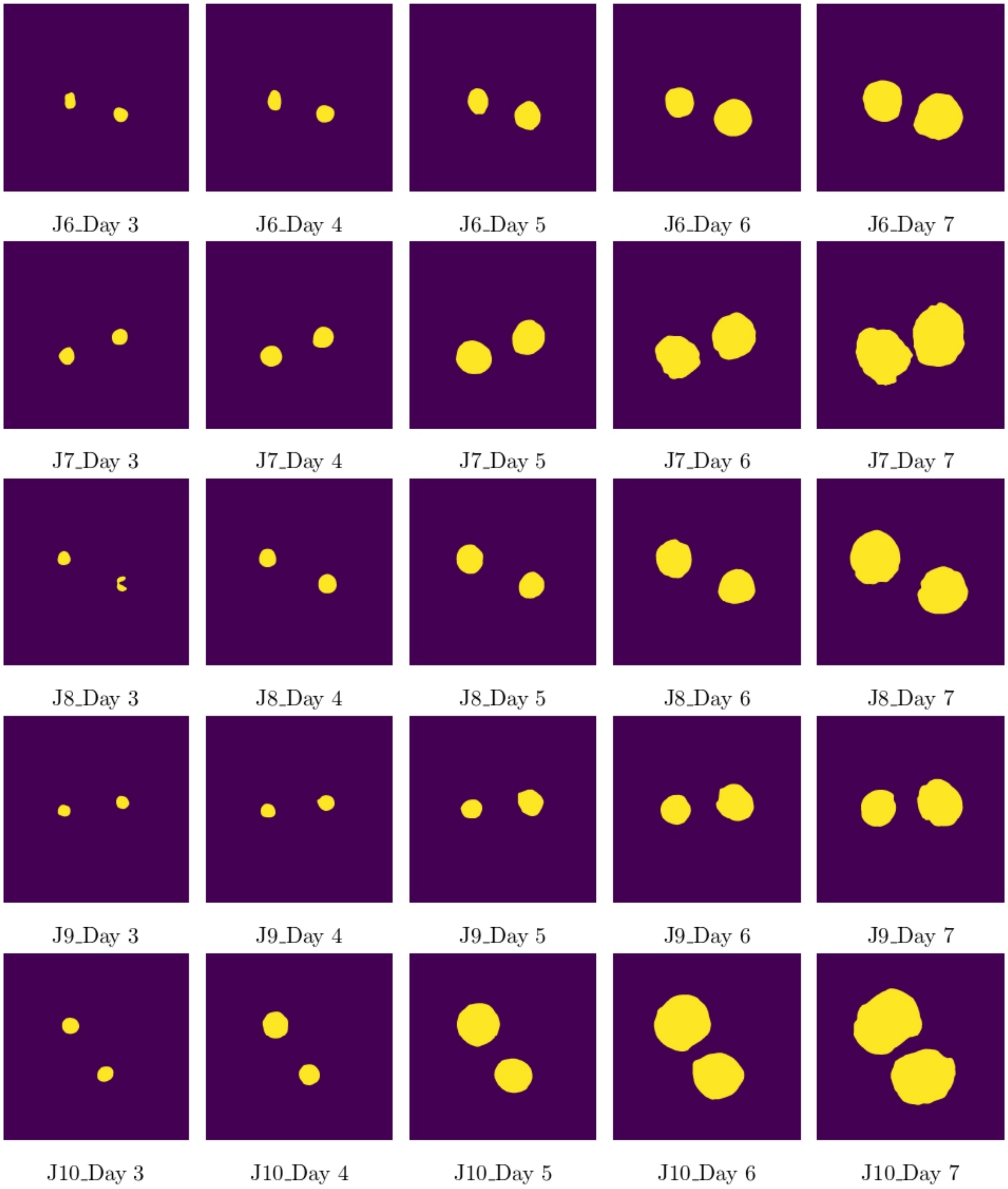


Figure 14: Binary images of lesion contours of *P. pinodes* on James cultivar *(continued)*

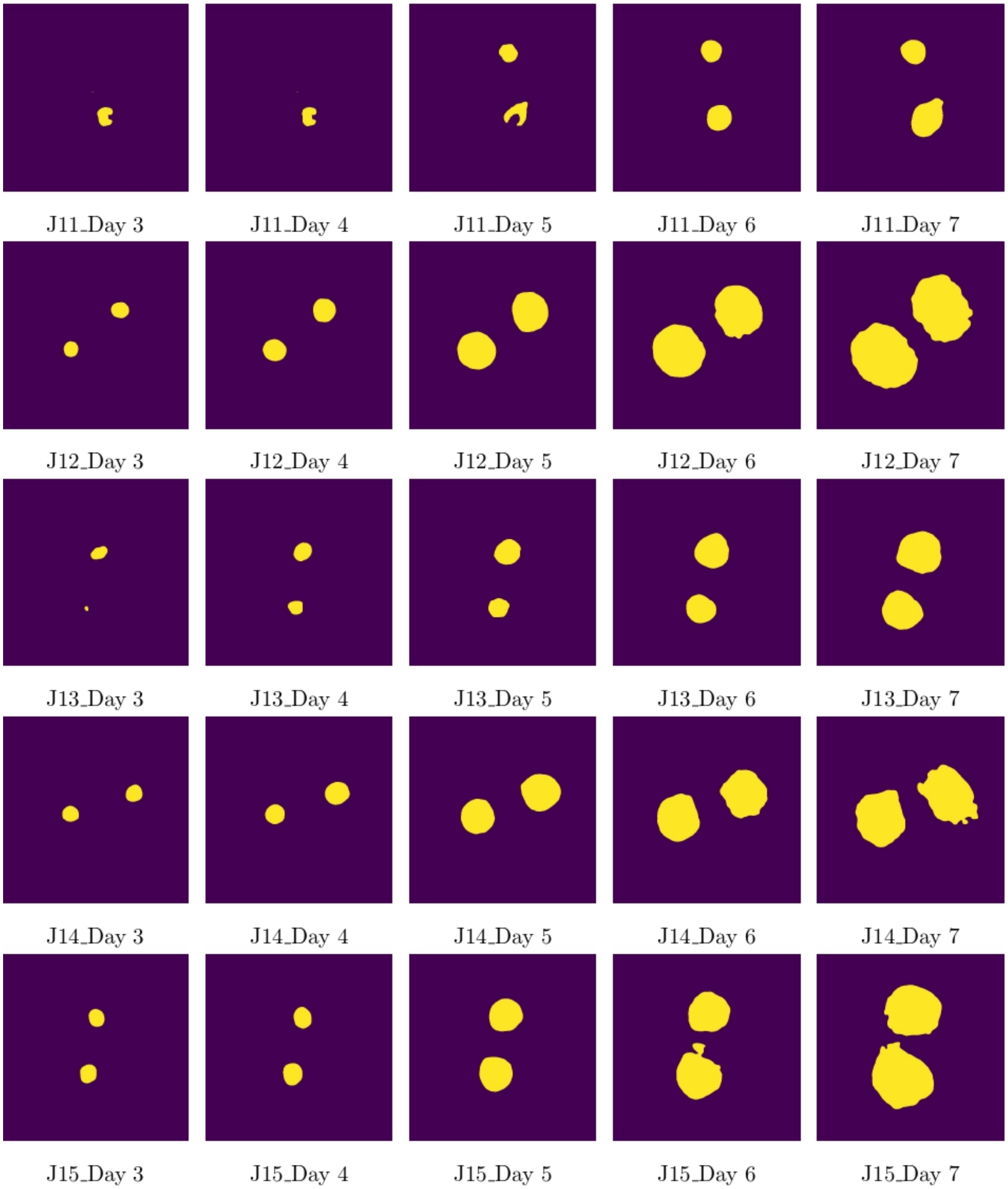


Figure 15: Binary images of lesion contours of *P. pinodes* on James cultivar *(continued)*

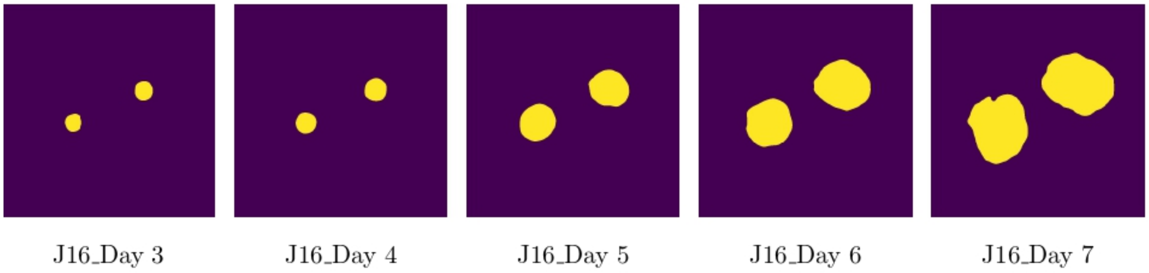


Figure 16: Binary images of lesion contours of *P. pinodes* on James cultivar *(continued)*

### S3. Level set simulation for leaf deformation

Figure 17 to Figure 20 and Figure 21 to Figure 24 show the comparison between the observed images of the leaf and level set solution of the thirty two (32) sets of images of Solara and James cultivar, respectively.

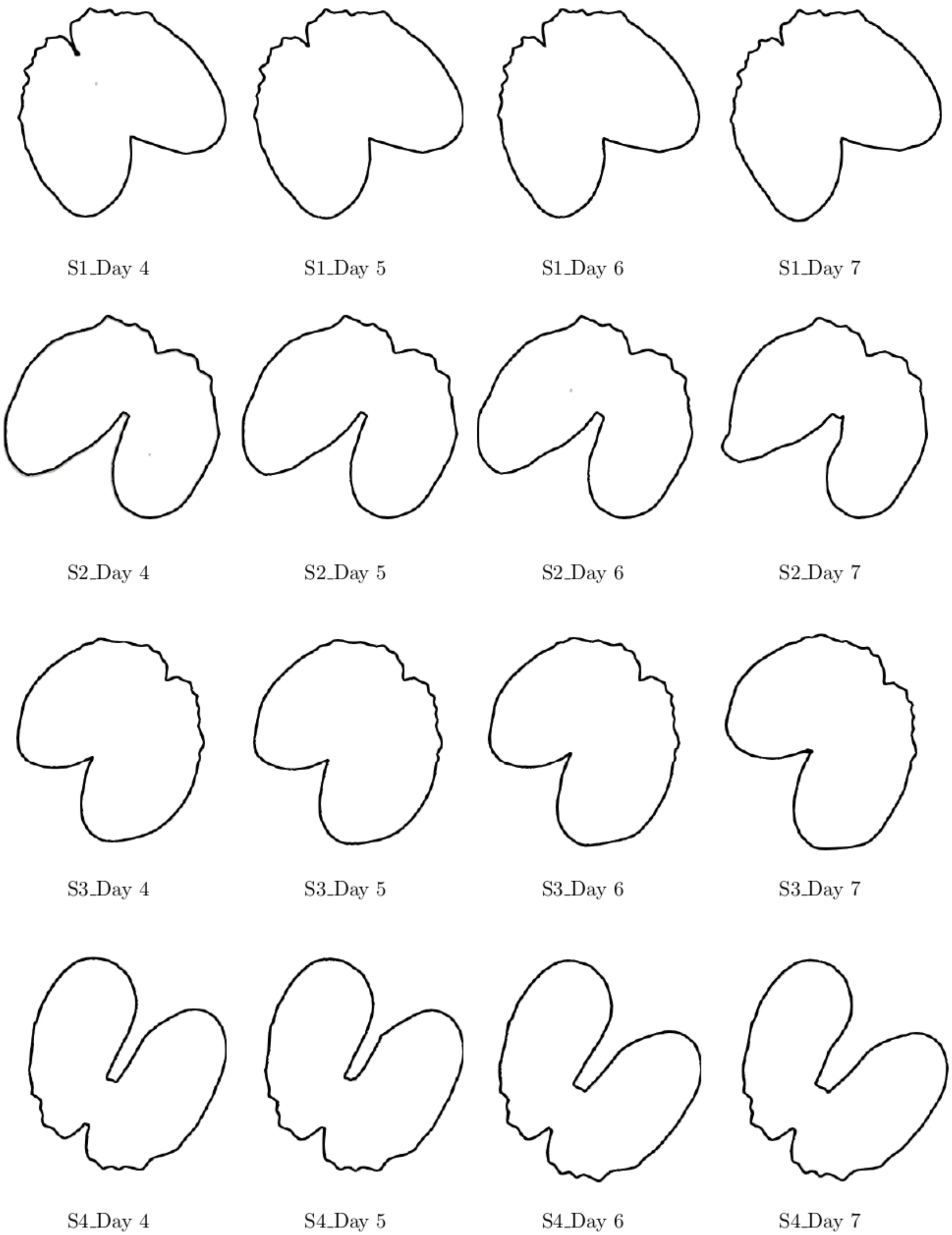


Figure 17: Comparison between observed images of leaf (gray contour) and level set solution (black contour) on Solara cultivar

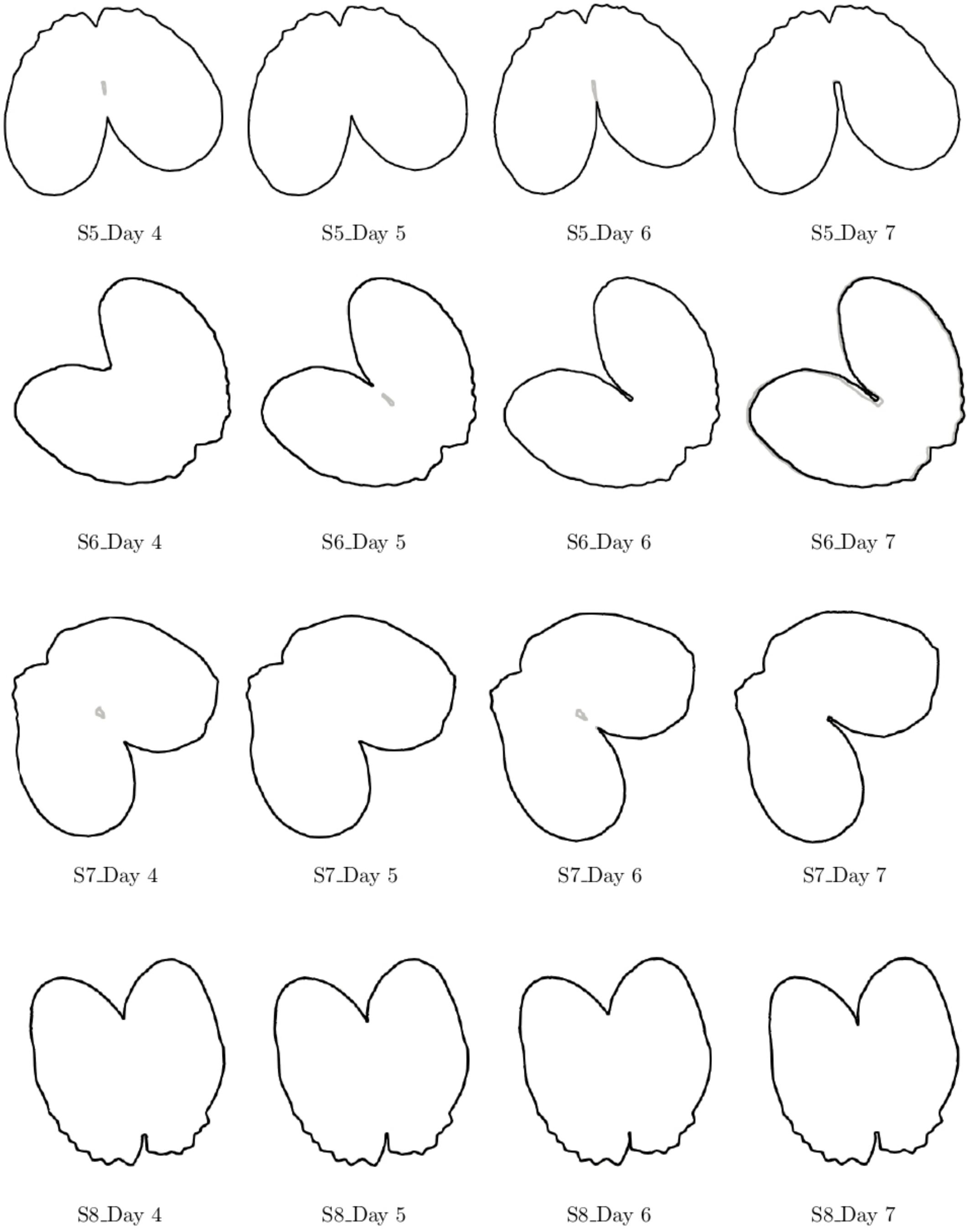


Figure 18: Comparison between observed images of leaf (gray contour) and level set solution (black contour) on Solara cultivar *(continued)*

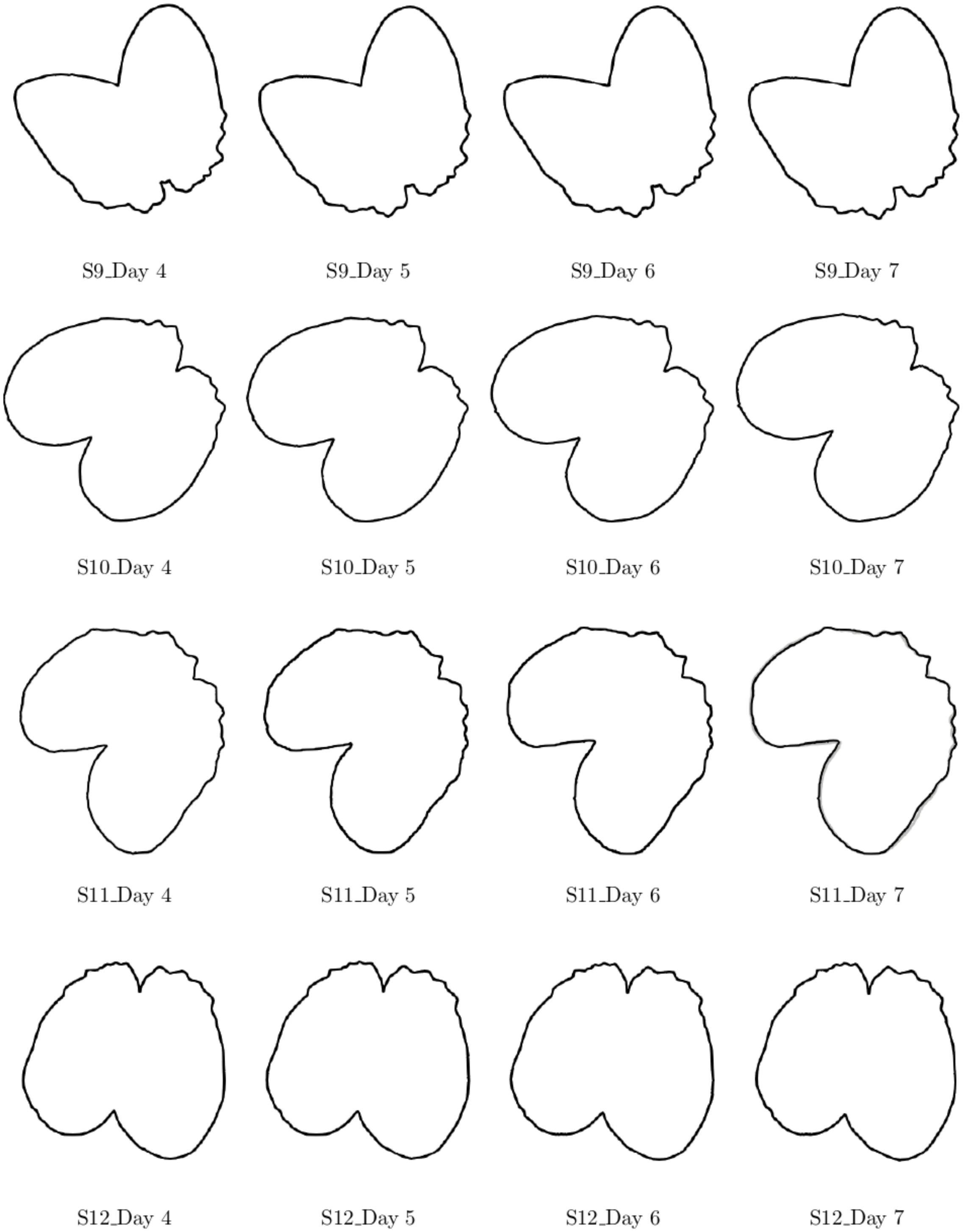


Figure 19: Comparison between observed images of leaf (gray contour) and level set solution (black contour) on Solara cultivar *(continued)*

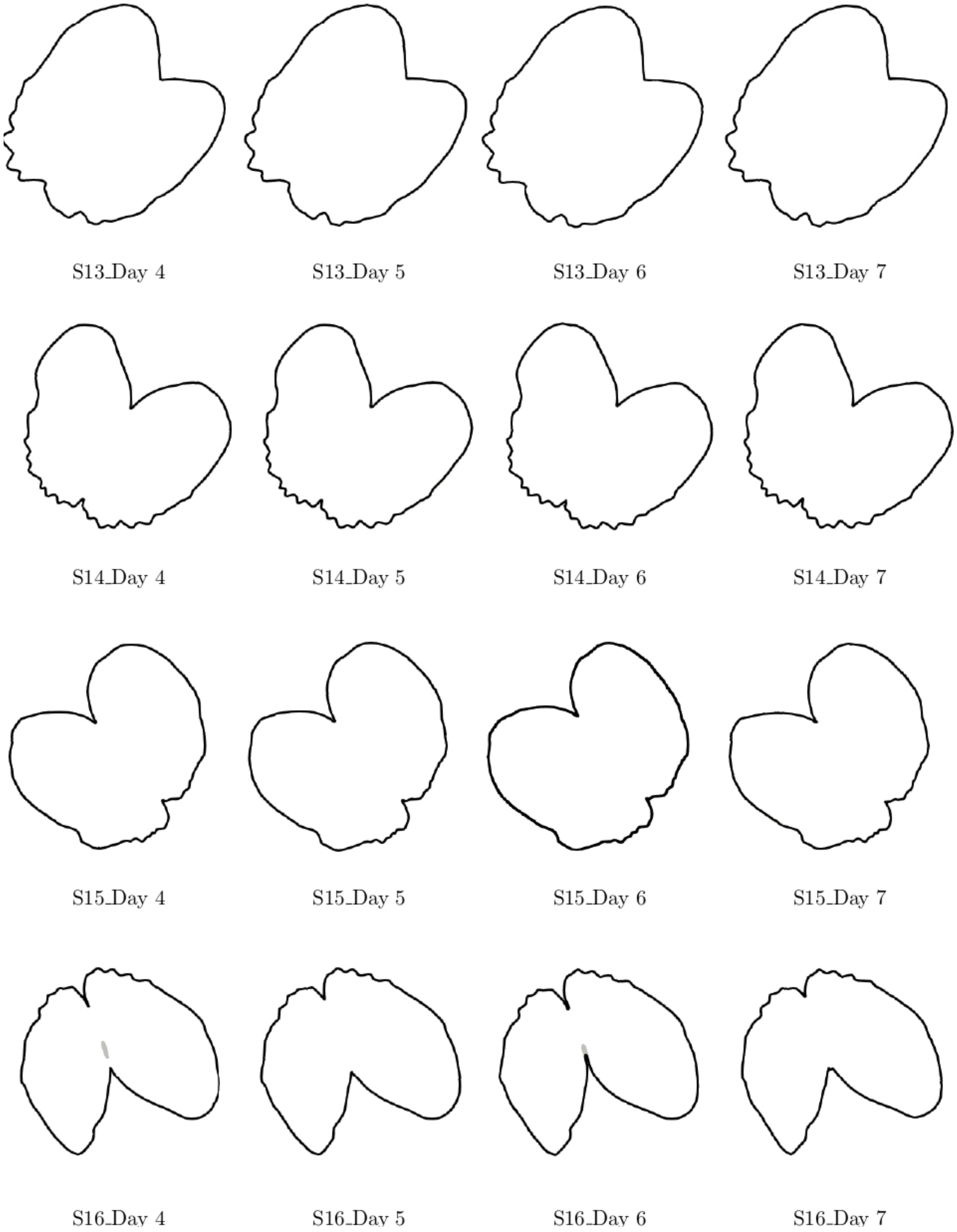


Figure 20: Comparison between observed images of leaf (gray contour) and level set solution (black contour) on Solara cultivar *(continued)*

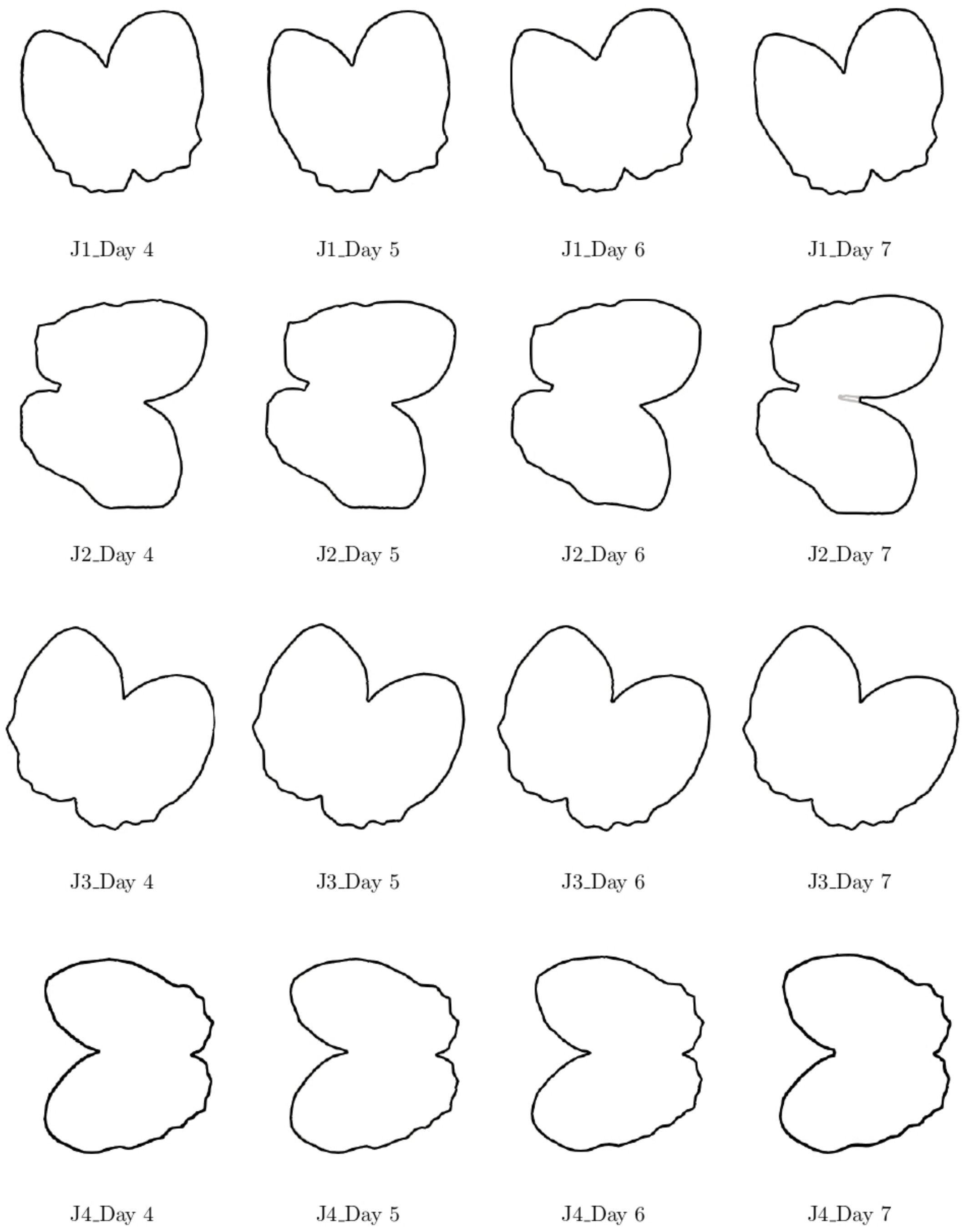


Figure 21: Comparison between observed images of leaf (gray contour) and level set solution (black contour) on James cultivar

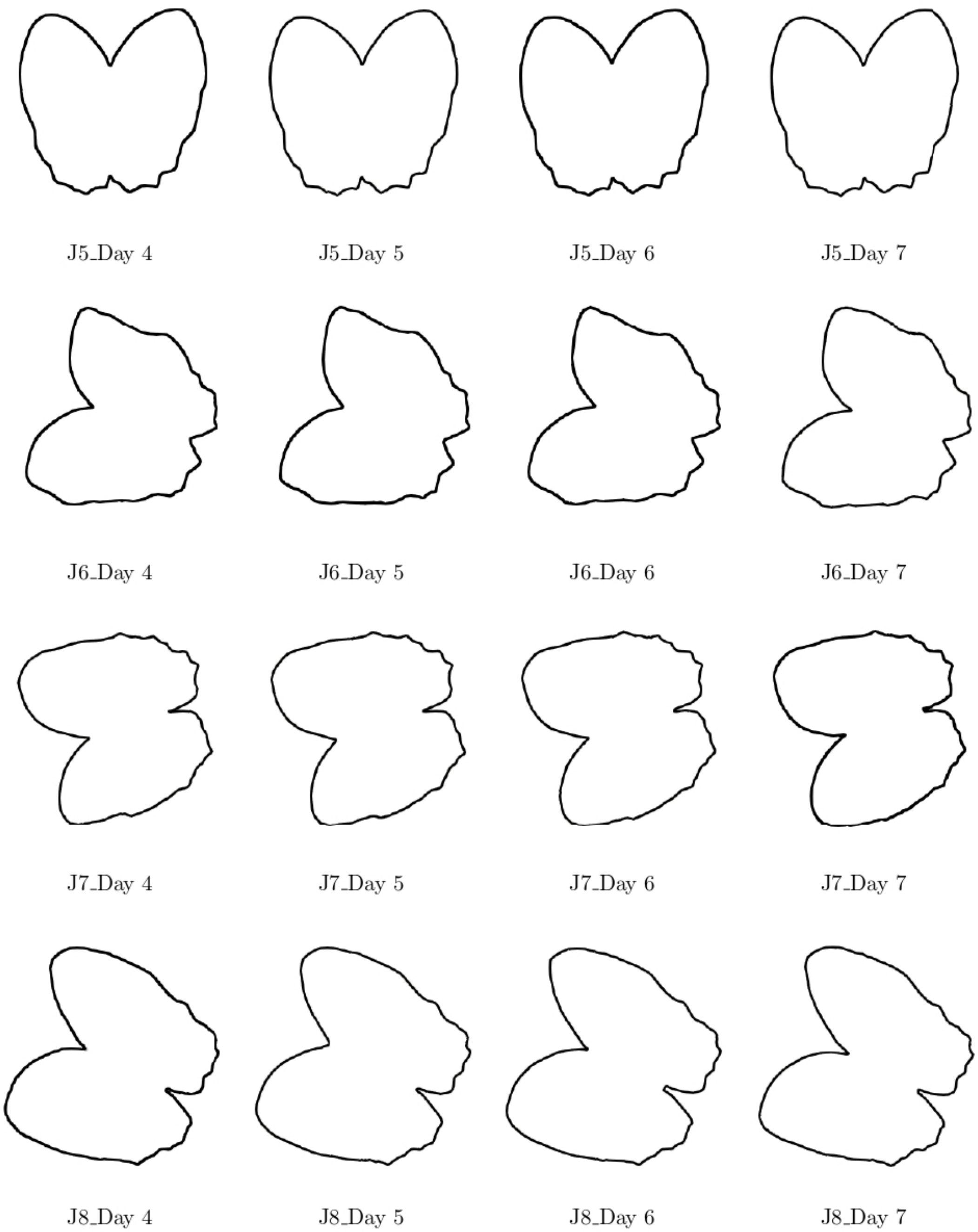


Figure 22: Comparison between observed images of leaf (gray contour) and level set solution (black contour) on James cultivar *(continued)*

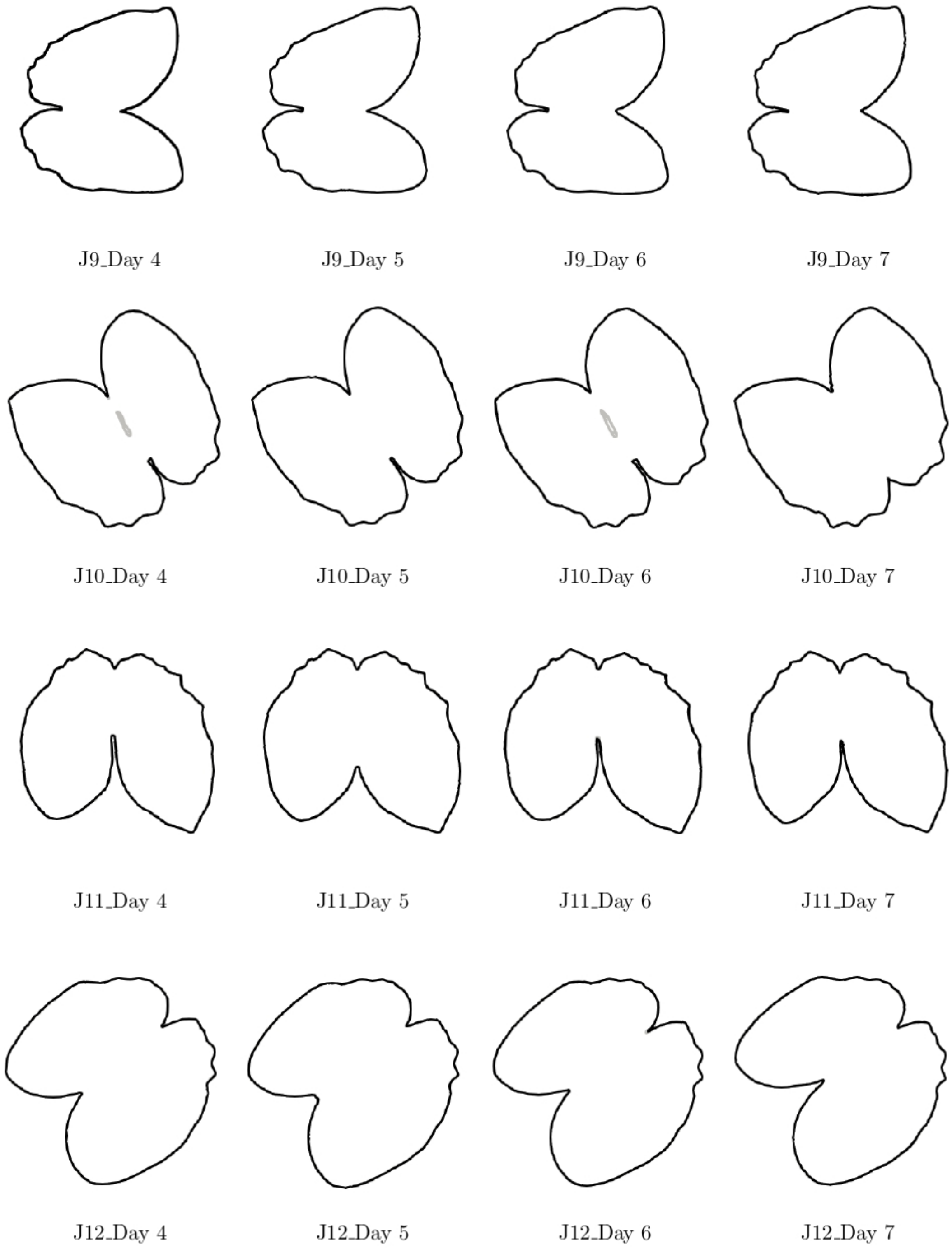


Figure 23: Comparison between observed images of leaf (gray contour) and level set solution (black contour) on James cultivar *(continued)*

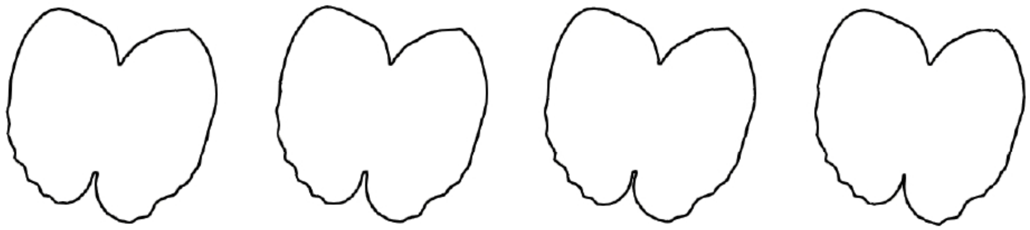

J13_Day 4 J13_Day 5 J13_Day 6 J13_Day 7

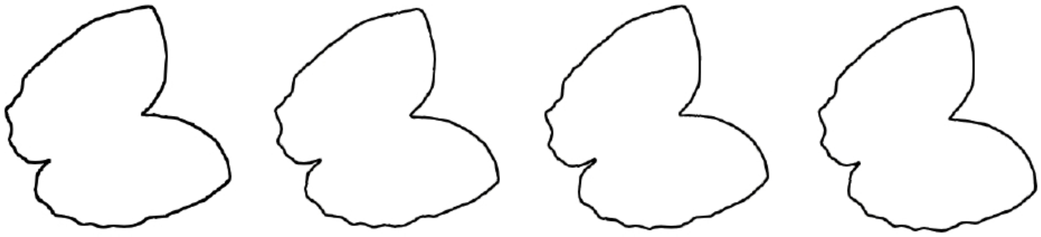

J14_Day 4 J14_Day 5 J14_Day 6 J14_Day 7

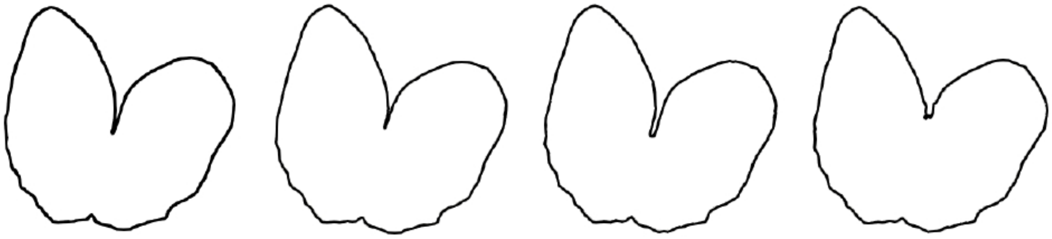

J15_Day 4 J15_Day 5 J15_Day 6 J15_Day 7

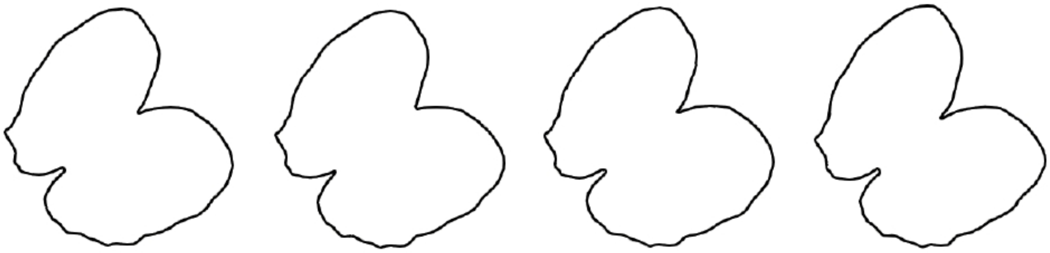

J16_Day 4 J16_Day 5 J16_Day 6 J16_Day 7

Figure 24: Comparison between observed images of leaf (gray contour) and level set solution (black contour) on James cultivar *(continued)*

### S4. Level set simulation for lesion growth

Figure 25 to Figure 28 and Figure 29 to Figure 32 show the comparison between the observed images of the leaf and level set solution of the thirty two (32) sets of images of Solara and James cultivar, respectively.

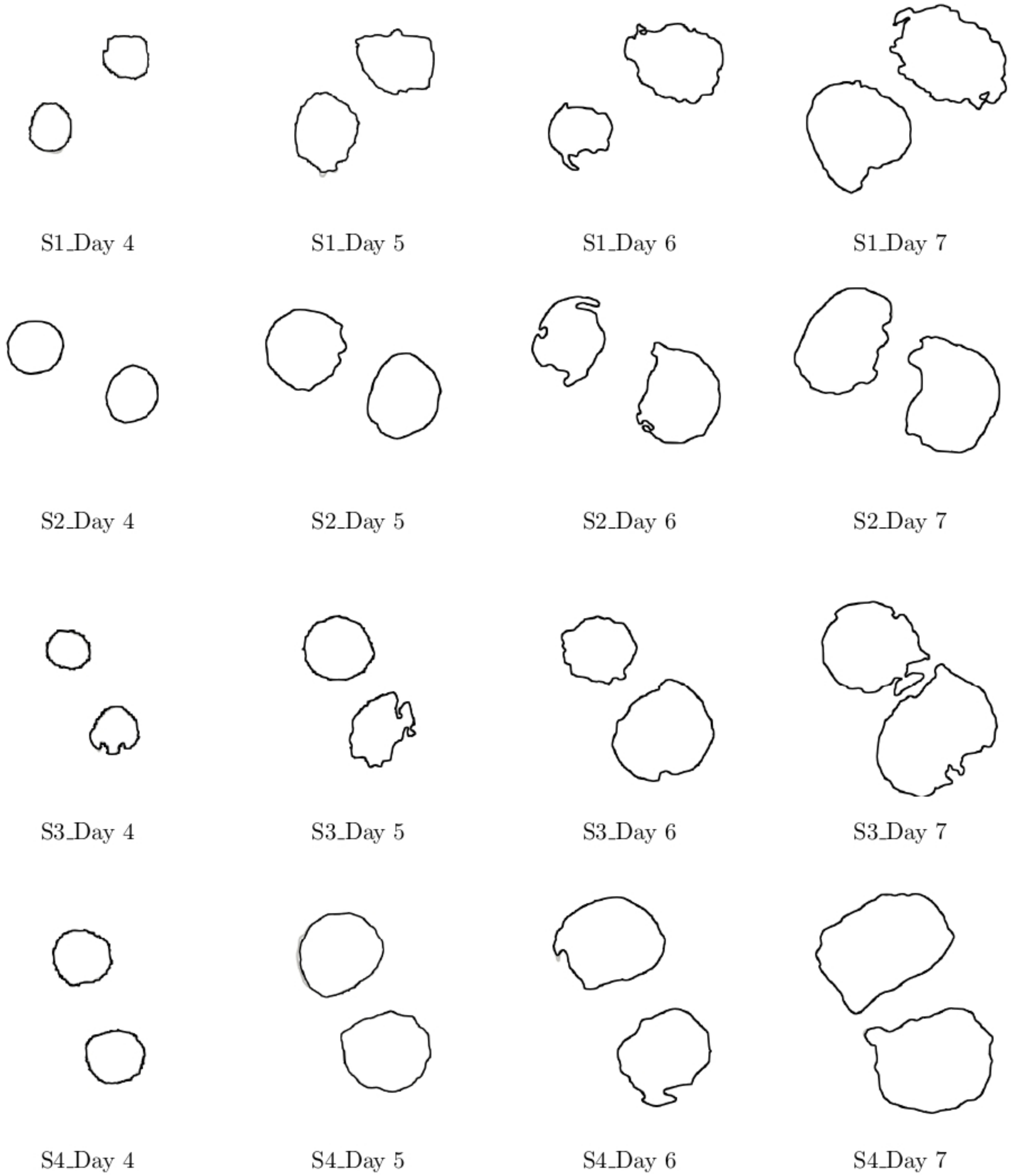


Figure 25: Comparison between observed images of lesion (gray contour) and level set solution (black contour) on Solara cultivar

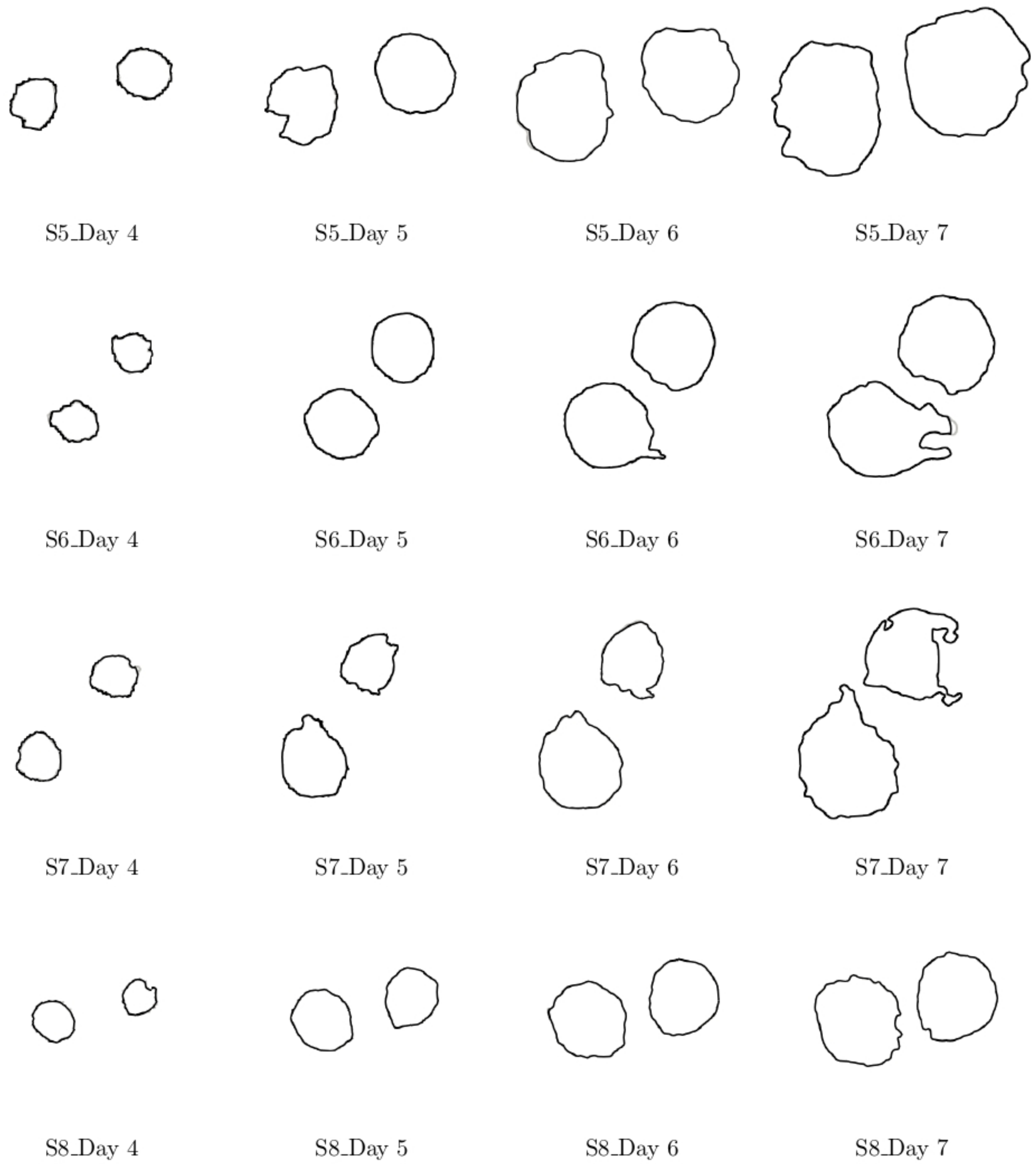


Figure 26: Comparison between observed images of lesion (gray contour) and level set solution (black contour) on Solara cultivar

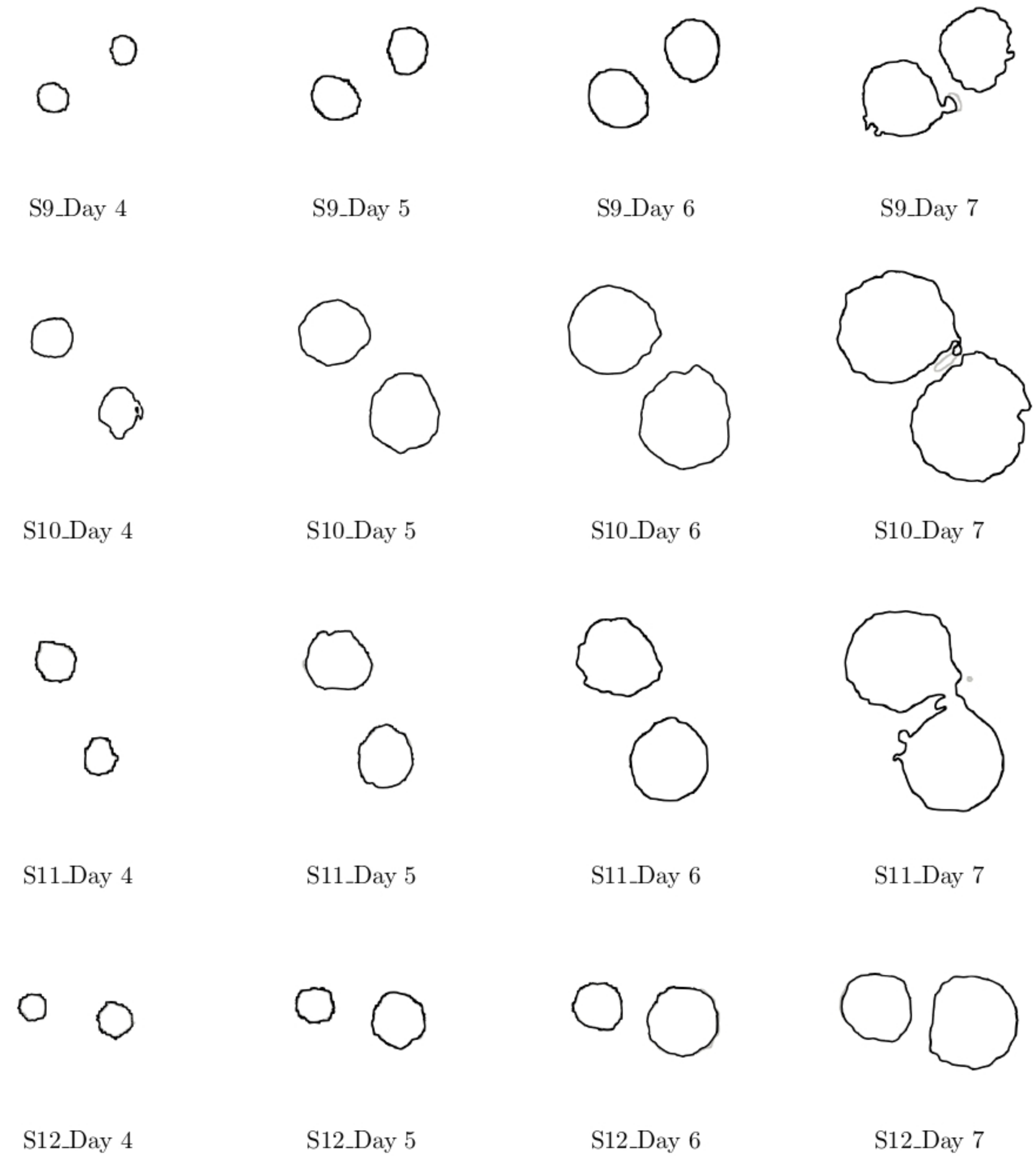


Figure 27: Comparison between observed images of lesion (gray contour) and level set solution (black contour) on Solara cultivar *(continued)*

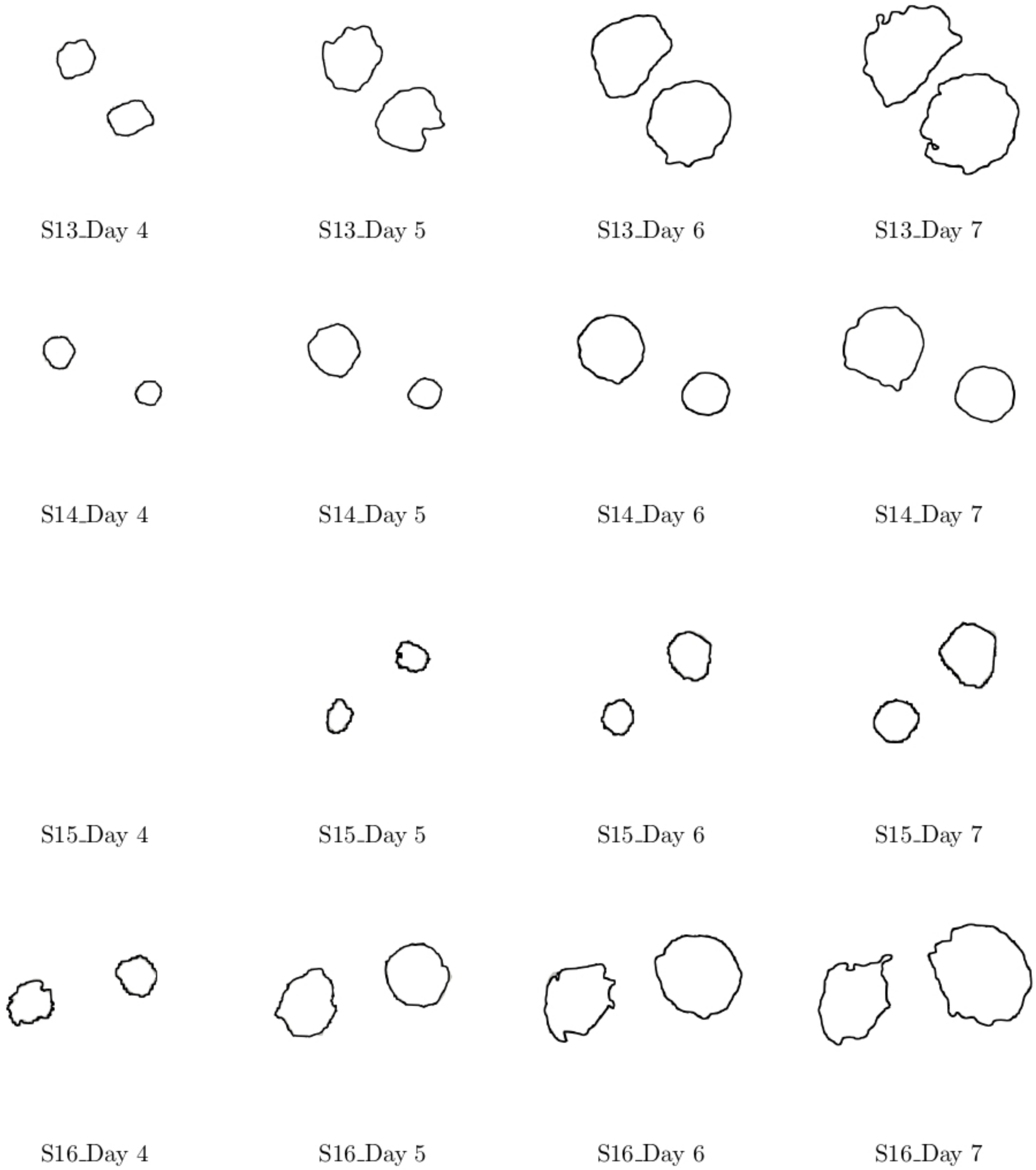


Figure 28: Comparison between observed images of lesion (gray contour) and level set solution (black contour) on Solara cultivar *(continued)*

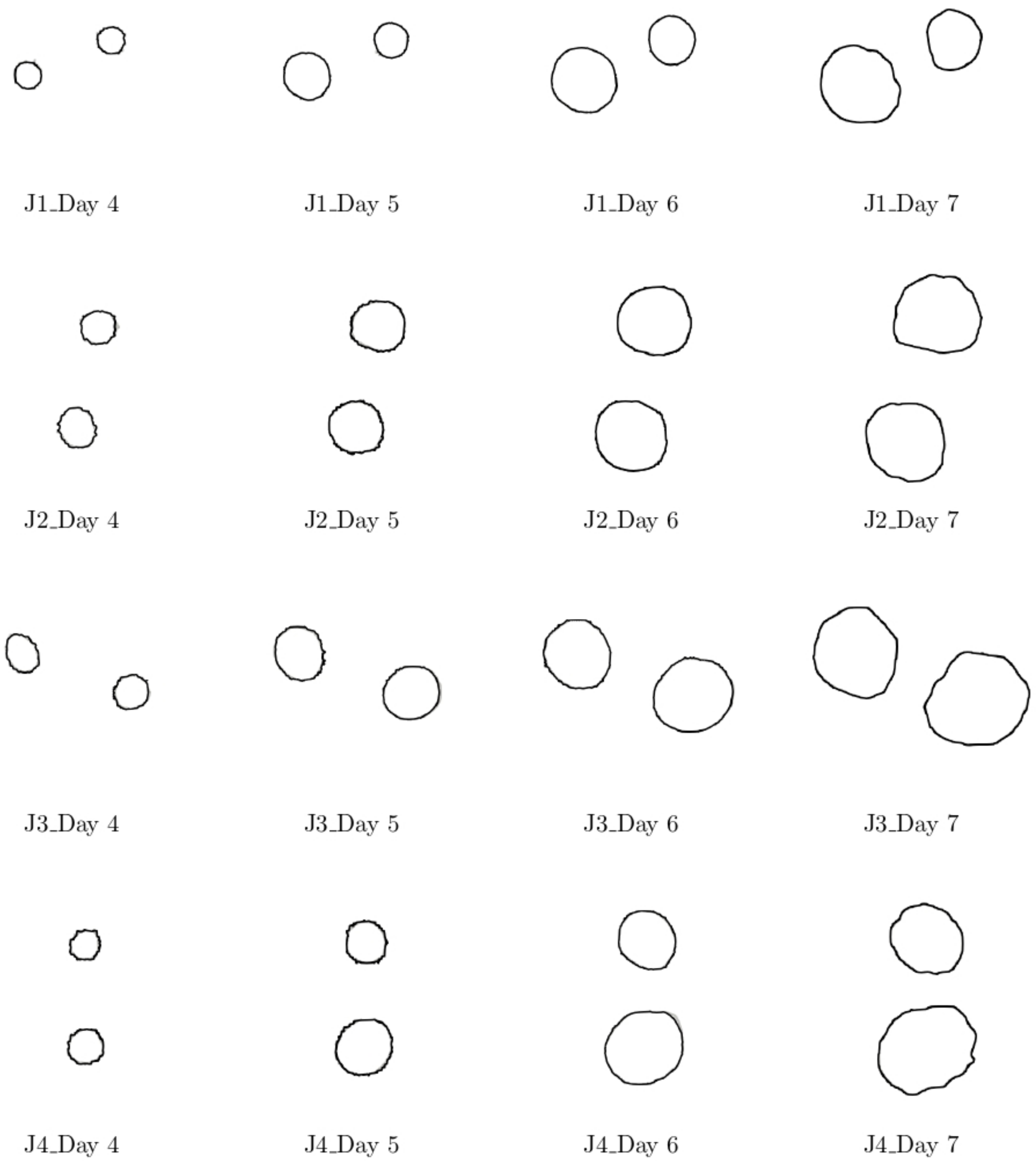


Figure 29: Comparison between observed images of lesion (gray contour) and level set solution (black contour) on James cultivar

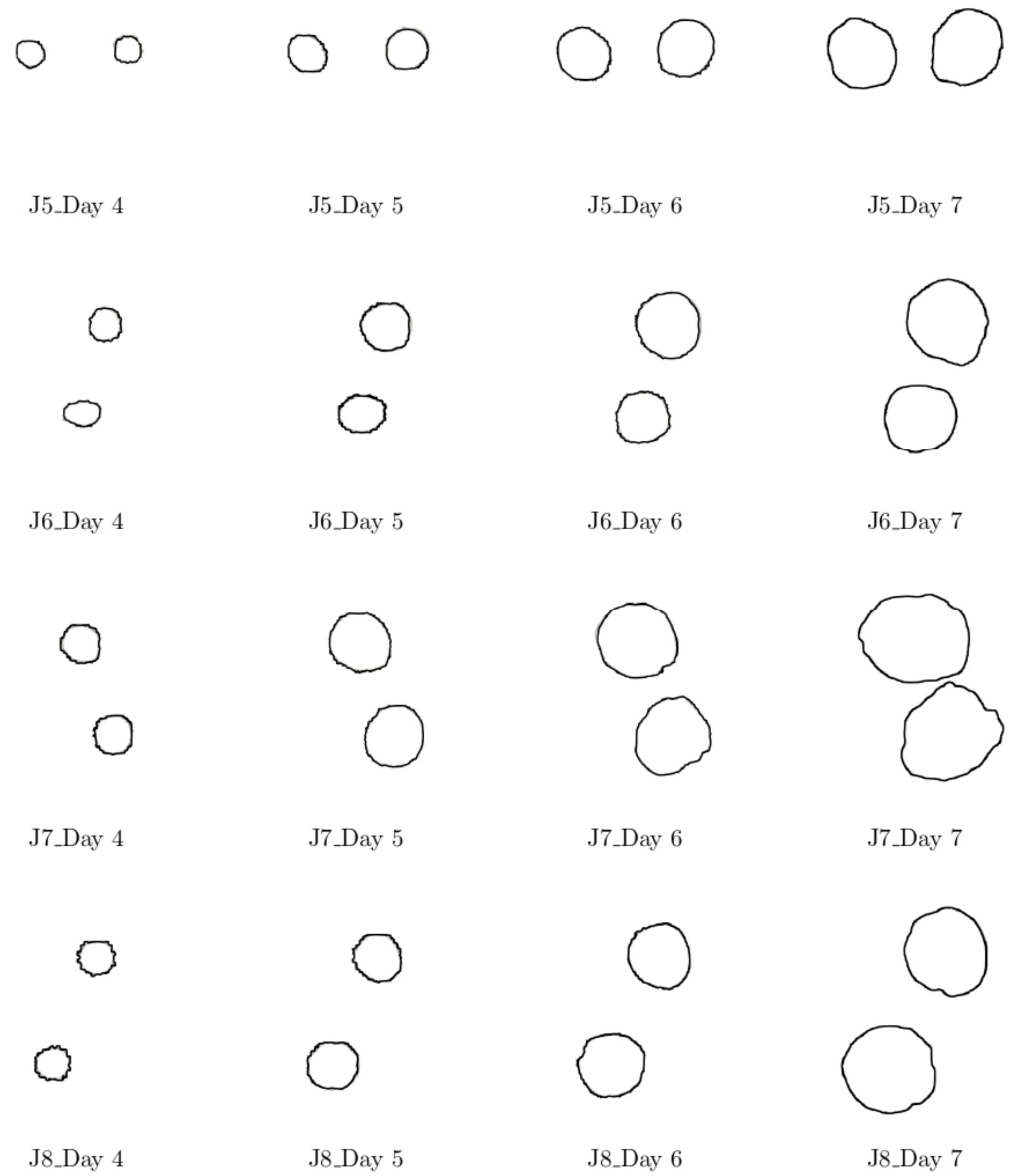


Figure 30: Comparison between observed images of lesion (gray contour) and level set solution (black contour) on James cultivar *(continued)*

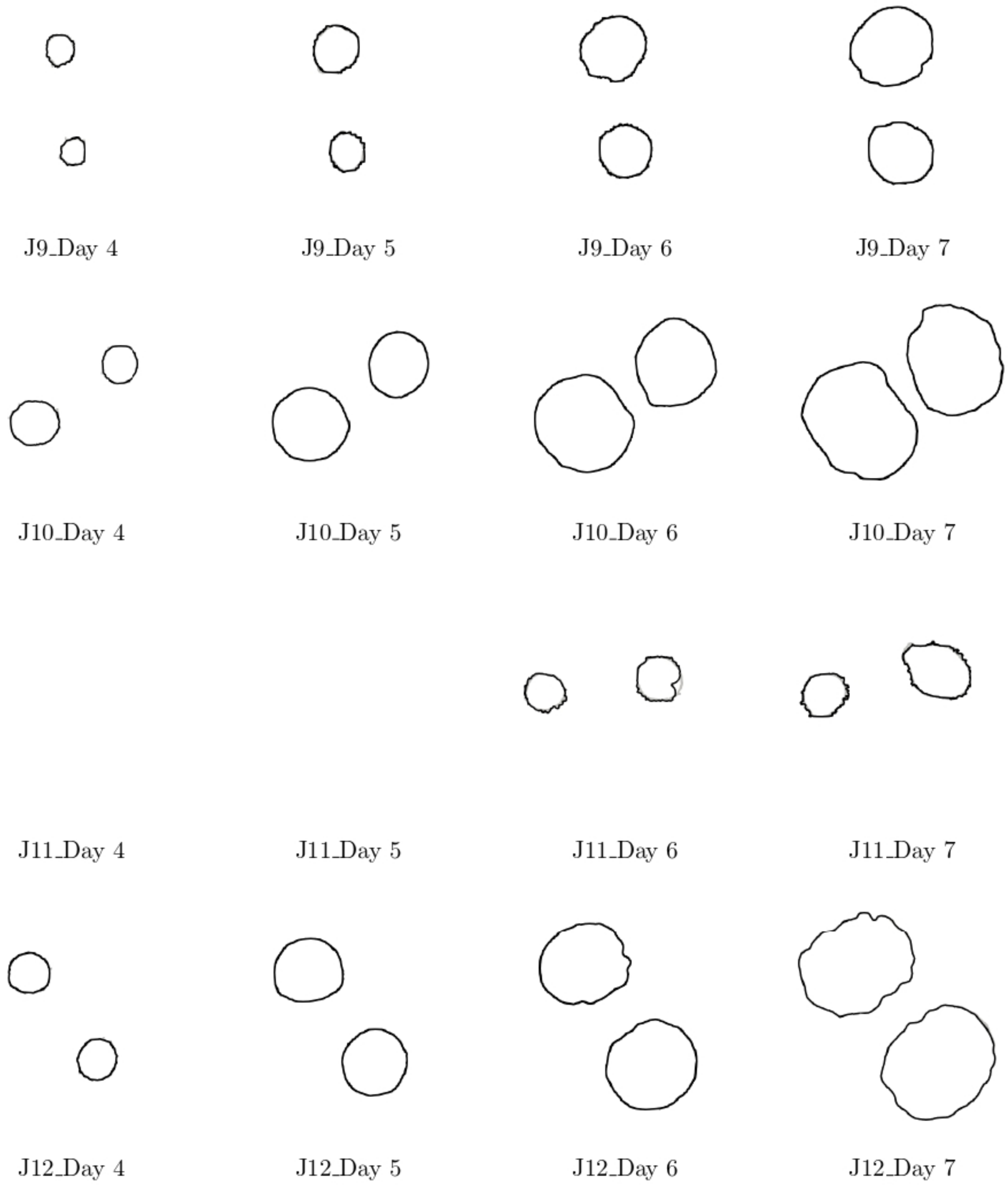


Figure 31: Comparison between observed images of lesion (gray contour) and level set solution (black contour) on James cultivar *(continued)*

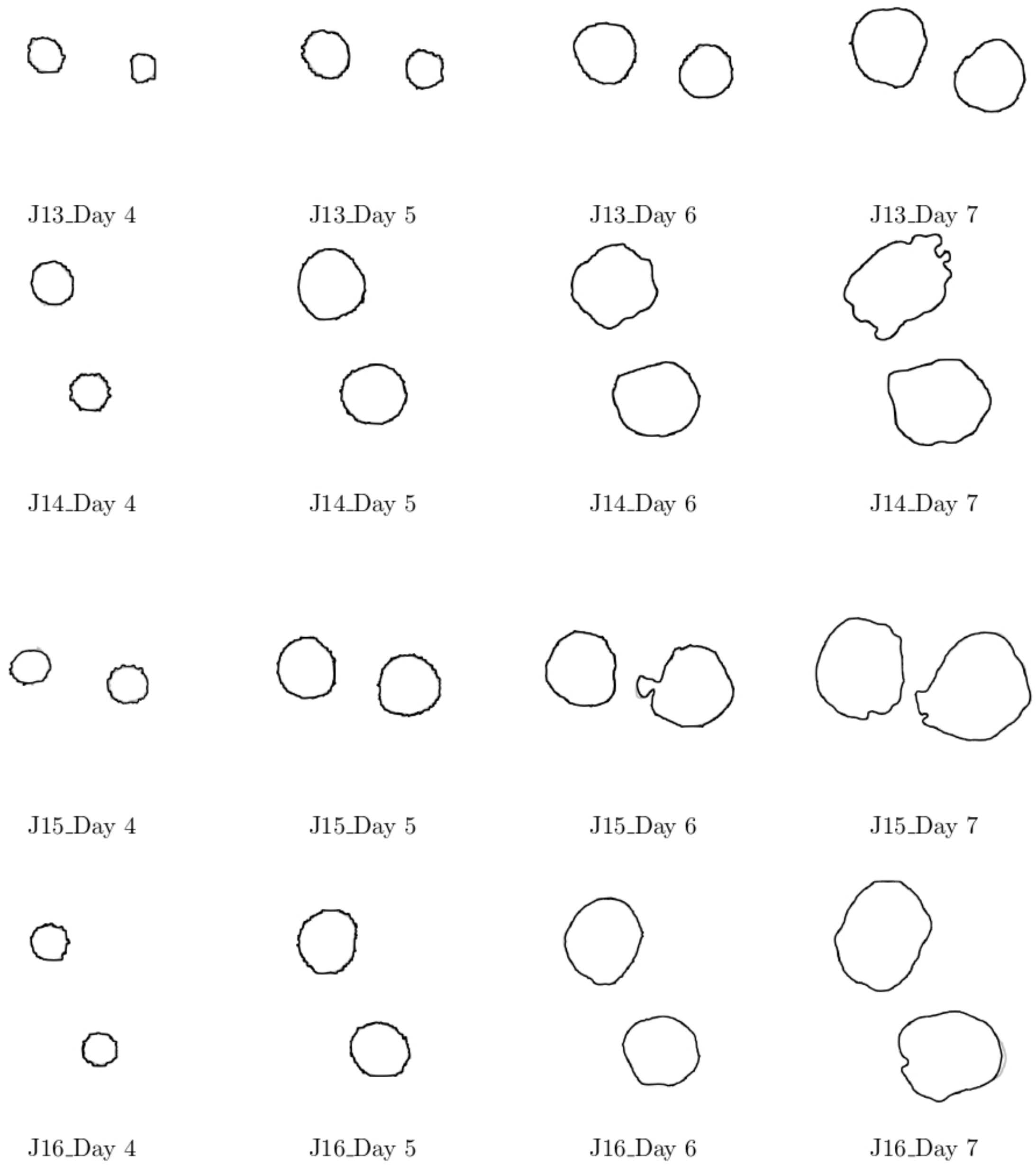


Figure 32: Comparison between observed images of lesion (gray contour) and level set solution (black contour) on James cultivar *(continued)*

### S5. Jaccard Similarity Index

*S5.1. Jaccard similarity index for leaf deformation*

| | Cultivar | Day 4 | Day 5 | Day 6 | Day 7 |
|---|---|---|---|---|---|
| 1 | Solara | 0.986745 | 0.993922 | 0.950042 | 0.966965 |
| 2 | Solara | 0.995167 | 0.985083 | 0.965043 | 0.869911 |
| 3 | Solara | 0.997070 | 0.988215 | 0.986197 | 0.952102 |
| 4 | Solara | 0.990784 | 0.983420 | 0.895736 | 0.876002 |
| 5 | Solara | 0.995645 | 0.993156 | 0.969656 | 0.958211 |
| 6 | Solara | 0.993368 | 0.977257 | 0.968497 | 0.959387 |
| 7 | Solara | 0.989871 | 0.991626 | 0.925494 | 0.905109 |
| 8 | Solara | 0.993134 | 0.988984 | 0.976691 | 0.966843 |
| 9 | Solara | 0.999078 | 0.980407 | 0.968378 | 0.964084 |
| 10 | Solara | 0.992237 | 0.987180 | 0.983986 | 0.967654 |
| 11 | Solara | 0.989994 | 0.993079 | 0.990951 | 0.977941 |
| 12 | Solara | 0.995510 | 0.994689 | 0.987648 | 0.970597 |
| 13 | Solara | 0.987058 | 0.977661 | 0.967748 | 0.965864 |
| 14 | Solara | 0.991725 | 0.987185 | 0.974867 | 0.970454 |
| 15 | Solara | 0.984659 | 0.988446 | 0.977146 | 0.967800 |
| 16 | Solara | 0.994049 | 0.993189 | 0.968609 | 0.952355 |
| 1 | James | 0.992807 | 0.988775 | 0.987844 | 0.973681 |
| 2 | James | 0.991895 | 0.974720 | 0.963688 | 0.950490 |
| 3 | James | 0.999583 | 0.989497 | 0.988956 | 0.984288 |
| 4 | James | 0.998781 | 0.995001 | 0.978827 | 0.972207 |
| 5 | James | 0.996994 | 0.987610 | 0.984931 | 0.976967 |
| 6 | James | 0.988215 | 0.979742 | 0.975442 | 0.982385 |
| 7 | James | 0.984124 | 0.988836 | 0.988064 | 0.980977 |
| 8 | James | 0.996126 | 0.993050 | 0.977637 | 0.956539 |
| 9 | James | 0.983441 | 0.977765 | 0.968194 | 0.979560 |
| 10 | James | 0.996557 | 0.986059 | 0.981817 | 0.973671 |
| 11 | James | 0.971373 | 0.972574 | 0.960168 | 0.942190 |
| 12 | James | 0.998193 | 0.988946 | 0.968135 | 0.954874 |
| 13 | James | 0.992404 | 0.995918 | 0.983802 | 0.978234 |
| 14 | James | 0.999811 | 0.974538 | 0.962186 | 0.938107 |
| 15 | James | 0.998252 | 0.978519 | 0.968198 | 0.959031 |
| 16 | James | 0.998416 | 0.983658 | 0.984456 | 0.959350 |

Table 1: Jaccard similarity index for leaf deformation of Solara and James cultivar between image data and level set solution

*S5.2. Jaccard similarity index for lesion growth*

| | Cultivar | Day 4 | Day 5 | Day 6 | Day 7 |
|---|---|---|---|---|---|
| 1 | Solara | 0.998514 | 0.995469 | 0.807326 | 0.993978 |
| 2 | Solara | 0.994726 | 0.998304 | 0.872436 | 0.979549 |
| 3 | Solara | 0.998559 | 0.973090 | 0.960197 | 0.988608 |
| 4 | Solara | 0.994268 | 0.992468 | 0.921997 | 0.985104 |
| 5 | Solara | 0.995651 | 0.996906 | 0.999961 | 0.999798 |
| 6 | Solara | 0.994889 | 0.994592 | 0.999411 | 0.987458 |
| 7 | Solara | 0.997125 | 0.909040 | 0.933099 | 0.984323 |
| 8 | Solara | 0.997474 | 0.993732 | 0.997693 | 0.999148 |
| 9 | Solara | 0.979100 | 0.998674 | 0.996033 | 0.997645 |
| 10 | Solara | 0.998370 | 0.9946540 | 0.999560 | 0.994020 |
| 11 | Solara | 0.995617 | 0.993137 | 0.997595 | 0.999086 |
| 12 | Solara | 0.981378 | 0.997245 | 0.994760 | 0.997742 |
| 13 | Solara | 0.997040 | 0.991921 | 0.996550 | 0.998453 |
| 14 | Solara | 0.982197 | 0.998254 | 0.995253 | 0.990214 |
| 15 | Solara | no lesion detected | 0.972008 | 0.996746 | 0.992005 |
| 16 | Solara | 0.998721 | 0.992323 | 0.964802 | 0.966113 |
| 1 | James | 0.956969 | 0.997998 | 0.994285 | 0.985800 |
| 2 | James | 0.975506 | 0.990782 | 0.991651 | 0.967335 |
| 3 | James | 0.993140 | 0.994200 | 0.994076 | 0.999211 |
| 4 | James | 0.986941 | 0.995623 | 0.993695 | 0.998039 |
| 5 | James | 0.963491 | 0.998519 | 0.994674 | 0.995079 |
| 6 | James | 0.980464 | 0.994814 | 0.994390 | 0.997062 |
| 7 | James | 0.998200 | 0.992730 | 0.996826 | 0.999912 |
| 8 | James | 0.987448 | 0.996142 | 0.989307 | 0.998995 |
| 9 | James | 0.987948 | 0.997820 | 0.992196 | 0.995849 |
| 10 | James | 0.995969 | 0.993969 | 0.999189 | 0.999961 |
| 11 | James | no lesion detected | no lesion detected | 0.928256 | 0.967641 |
| 12 | James | 0.997829 | 0.992747 | 0.998502 | 0.999388 |
| 13 | James | 0.963934 | 0.999277 | 0.991314 | 0.992566 |
| 14 | James | 0.995804 | 0.990791 | 0.996063 | 0.999233 |
| 15 | James | 0.981857 | 0.989547 | 0.994170 | 0.989179 |
| 16 | James | 0.989340 | 0.990223 | 0.995870 | 0.999810 |

Table 2: Jaccard similarity index for leaf deformation of Solara and James cultivar between image data and level set solution

### S6. Relative error

*S6.1. Relative error for the simulation of the leaf deformation*

| | Cultivar | Day 4 | Day 5 | Day 6 | Day 7 |
|---|---|---|---|---|---|
| 1 | Solara | 0.005300 | 0.020607 | 0.015745 | 0.009524 |
| 2 | Solara | 0.008189 | 0.241431 | 0.020247 | 0.005181 |
| 3 | Solara | 0.010925 | 0.008265 | 0.013951 | 0.003837 |
| 4 | Solara | 0.005027 | 0.036396 | 0.000996 | 0.014465 |
| 5 | Solara | 0.004387 | 0.055504 | 0.018157 | 0.004836 |
| 6 | Solara | 0.003082 | 0.034297 | 0.010946 | 0.024663 |
| 7 | Solara | 0.028319 | 0.043361 | 0.004347 | 0.007497 |
| 8 | Solara | 0.001591 | 0.056904 | 0.016707 | 0.002928 |
| 9 | Solara | 0.008434 | 0.004472 | 0.004042 | 0.005105 |
| 10 | Solara | 0.008087 | 0.033913 | 0.003971 | 0.002713 |
| 11 | Solara | 0.012219 | 0.015499 | 0.095713 | 0.008904 |
| 12 | Solara | 0.001903 | 0.391872 | 0.053089 | 0.015529 |
| 13 | Solara | 0.006812 | 0.004511 | 0.004402 | 0.061806 |
| 14 | Solara | 0.002099 | 0.101425 | 0.003253 | 0.007486 |
| 15 | Solara | 0.004241 | 0.007402 | 0.007215 | 0.008593 |
| 16 | Solara | 0.015447 | 0.168339 | 0.016279 | 0.004114 |
| 17 | James | 0.013349 | 0.499622 | 0.012749 | 0.003479 |
| 18 | James | 0.016741 | 0.011411 | 0.021376 | 0.026647 |
| 19 | James | 0.019283 | 0.012994 | 0.027949 | 0.018783 |
| 20 | James | 0.008613 | 0.143796 | 0.009372 | 0.036275 |
| 21 | James | 0.020149 | 0.013568 | 0.196045 | 0.087505 |
| 22 | James | 0.014360 | 0.027867 | 0.026059 | 0.008748 |
| 23 | James | 0.025602 | 0.021796 | 0.050866 | 0.050393 |
| 24 | James | 0.005933 | 1.756606 | 0.047670 | 0.132451 |
| 25 | James | 0.012446 | 0.018190 | 0.022773 | 0.010870 |
| 26 | James | 0.047621 | 0.013081 | 0.038240 | 0.019106 |
| 27 | James | 0.003952 | 0.016146 | 0.070310 | 0.014877 |
| 28 | James | 0.135641 | 0.041106 | 0.013565 | 0.004210 |
| 29 | James | 0.005298 | 0.424251 | 0.037720 | 0.067455 |
| 30 | James | 0.005660 | 0.025770 | 0.013715 | 0.005120 |
| 31 | James | 0.028690 | 0.008664 | 0.041067 | 0.007472 |
| 32 | James | 0.006730 | 0.029176 | 0.020007 | 0.011408 |

Table 3: Relative errors for leaf deformation of Solara and James cultivars between data image and level set solution

*S6.2. Daily mean relative errors for leaf deformation*

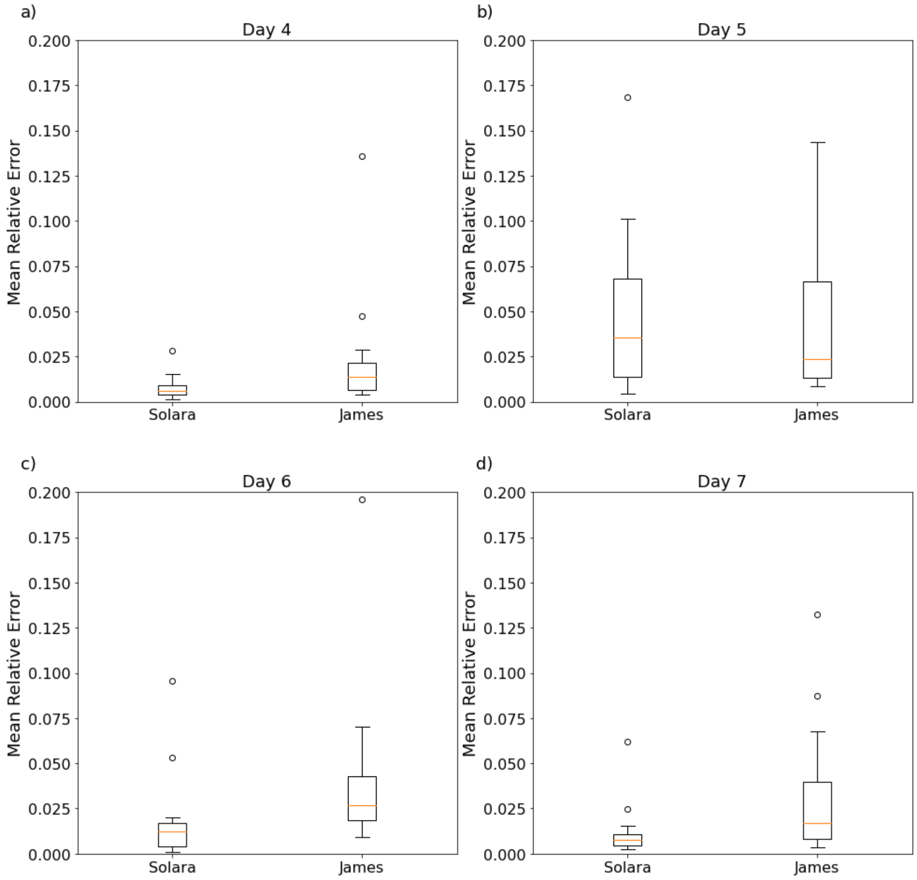


Figure 33: Boxplot for daily mean relative errors for leaf deformation of Solara and James cultivars

| Day | Cultivar | Mean Relative Error | Minimum | Maximum | Outliers |
|---|---|---|---|---|---|
| Day 4 | Solara | 0.007878875 | 0.001591 | 0.00395125 | 0.028319 |
| | James | 0.02312925 | 0.015447 | 0.00905675 | 0.047621 |
| | | | | | 0.135641 |
| Day 5 | Solara | 0.076512375 | 0.004472 | 0.0136905 | 0.241431 |
| | | | | | 0.391872 |
| | | | | | 0.168339 |
| | James | 0.19150275 | 0.101425 | 0.06803425 | 0.499622 |
| | | | | | 1.756606 |
| | | | | | 0.424251 |
| Day 6 | Solara | 0.01806625 | 0.000996 | 0.00427075 | 0.095713 |
| | | | | | 0.053089 |
| | James | 0.0405926875 | 0.020247 | 0.0170695 | 0.196045 |
| Day 7 | Solara | 0.0116988125 | 0.002713 | 0.0046555 | 0.024663 |
| | | | | | 0.061806 |
| | James | 0.0315499375 | 0.015529 | 0.01075925 | 0.087505 |
| | | | | | 0.132451 |

Table 4: Daily mean relative errors for leaf deformation of Solara and James cultivars

*S6.3. Relative error for the simulation of lesion growth*

| | Cultivar | Day 4 | Day 5 | Day 6 | Day 7 |
|---|---|---|---|---|---|
| 1 | Solara | 0.002849 | 0.000915 | 0.002091 | 0.005146 |
| 2 | Solara | 0.001131 | 0.000906 | 0.011829 | 0.001636 |
| 3 | Solara | 0.004400 | 0.000931 | 0.016287 | 0.002769 |
| 4 | Solara | 0.001796 | 0.000908 | 0.001853 | 0.001436 |
| 5 | Solara | 0.004638 | 0.000923 | 0.000909 | 0.001167 |
| 6 | Solara | 0.003439 | 0.000959 | 0.003255 | 0.006538 |
| 7 | Solara | 0.003163 | 0.001851 | 0.000987 | 0.005097 |
| 8 | Solara | 0.010995 | 0.000866 | 0.000724 | 0.000820 |
| 9 | Solara | 0.119205 | 0.001640 | 0.002532 | 0.004371 |
| 10 | Solara | 0.006516 | 0.000894 | 0.000993 | 0.002355 |
| 11 | Solara | 0.021507 | 0.000987 | 0.000986 | 0.328940 |
| 12 | Solara | 0.041229 | 0.004419 | 0.002771 | 0.000878 |
| 13 | Solara | 0.012499 | 0.000997 | 0.000939 | 0.001333 |
| 14 | Solara | 0.023383 | 0.134649 | 0.002123 | 0.002078 |
| 15 | Solara | no lesion detected | 0.028192 | 0.013992 | 0.002517 |
| 16 | Solara | 0.009382 | 0.001850 | 0.001145 | 0.005818 |
| 17 | James | 0.105642 | 0.027200 | 0.003846 | 0.002725 |
| 18 | James | 0.097874 | 0.003773 | 0.001827 | 0.001936 |
| 19 | James | 0.048844 | 0.003851 | 0.001623 | 0.000823 |
| 20 | James | 0.116723 | 0.005027 | 0.001584 | 0.001627 |
| 21 | James | 0.473685 | 0.007869 | 0.002129 | 0.000965 |
| 22 | James | 0.121596 | 0.004592 | 0.002589 | 0.000851 |
| 23 | James | 0.037729 | 0.000939 | 0.000982 | 0.000878 |
| 24 | James | 0.119502 | 0.005507 | 0.001655 | 0.000754 |
| 25 | James | 0.114708 | 0.011728 | 0.001526 | 0.001653 |
| 26 | James | 0.017912 | 0.001243 | 0.001397 | 0.001047 |
| 27 | James | no lesion detected | no lesion detected | 0.026698 | 0.033580 |
| 28 | James | 0.005718 | 0.000944 | 0.000860 | 0.000995 |
| 29 | James | 0.015047 | 0.008960 | 0.002711 | 0.001565 |
| 30 | James | 0.118358 | 0.001496 | 0.002641 | 0.001135 |
| 31 | James | 0.070462 | 0.002391 | 0.001700 | 0.000831 |
| 32 | James | 0.083149 | 0.001507 | 0.001484 | 0.000884 |

Table 5: Relative errors for lesion growth of Solara and James cultivars between data image and level set solution

*S6.4. Daily mean relative errors for lesion growth*

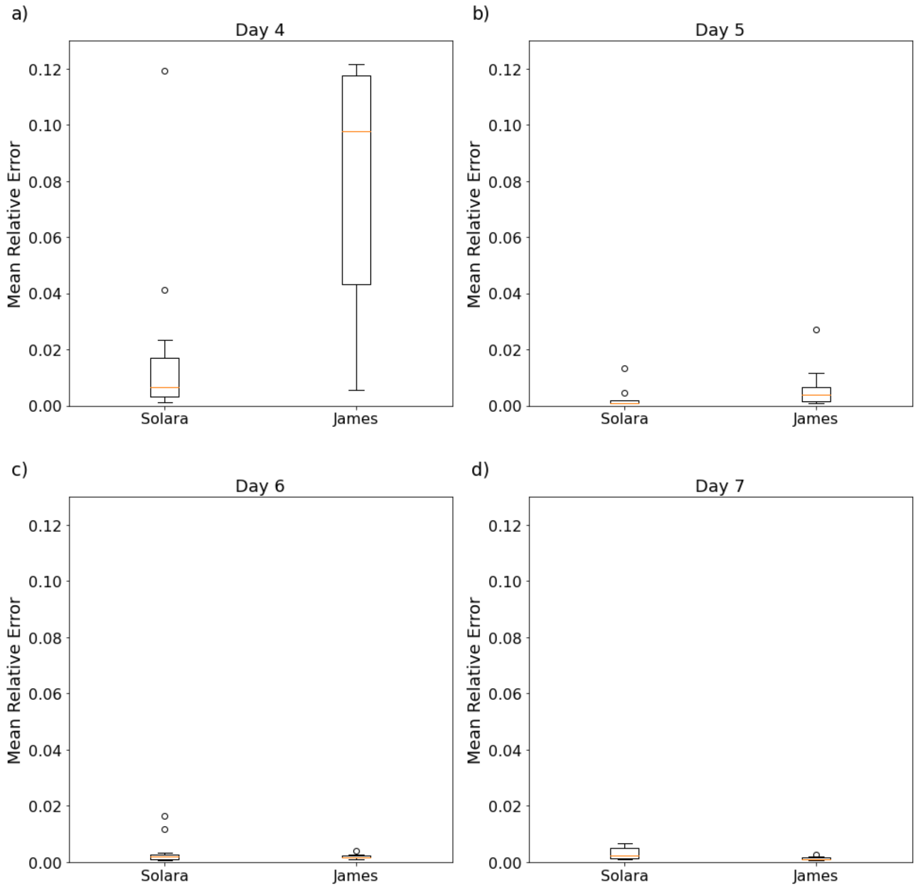


Figure 34: Boxplot for daily mean relative errors for lesion growth of Solara and James cultivars

| Day | Cultivar | Mean Relative Error | Minimum | Maximum | Outliers |
|---|---|---|---|---|---|
| Day 4 | Solara | 0.0177423 | 0.0011315 | 0.0033012 | 0.11920523 |
| | | | | | 0.04122872 |
| | James | 0.1031302 | 0.0233827 | 0.0170032 | 0.47368529 |
| Day 5 | Solara | 0.0021675 | 0.0008656 | 0.0009119 | 0.00441891 |
| | | | | | 0.0134649 |
| | James | 0.0058019 | 0.0018507 | 0.0017448 | 0.02720048 |
| Day 6 | Solara | 0.0032949 | 0.0007240 | 0.0009865 | 0.01182906 |
| | | | | | 0.0162866 |
| | James | 0.0019038 | 0.0032552 | 0.0026517 | 0.00384595 |
| Day 7 | Solara | 0.0246922 | 0.0008196 | 0.0013845 | 0.32894048 |
| | James | 0.0012448 | 0.0065376 | 0.0051217 | 0.00272523 |

Table 6: Daily mean relative errors for lesion growth of Solara and James cultivars
